# Cellular and circuit mechanisms underlying binocular vision

**DOI:** 10.1101/2024.03.11.584536

**Authors:** Suraj Honnuraiah, Helena Huang, William J. Ryan, Robin Broersen, William M. Connelly, Greg J. Stuart

## Abstract

How binocular visual information is combined at the level of single neurons in the brain is not fully understood. Here, we show in mice that callosal input from the opposite visual cortex (V1) plays a critical role in this process. *In vivo* we find this callosal projection carries ipsilateral eye information and synapses exclusively onto binocular neurons. Using the presence of callosal input to identify binocular neurons *in vitro,* at the cellular level we show that binocular neurons are less excitable than monocular neurons due to high expression of Kv1 potassium channels. At the circuit level we find that only monocular neurons send callosal projections to the opposite V1, whereas binocular neurons do not. Finally, using dual-colour optogenetics we show that most binocular and monocular neurons receive direct input from the thalamus. In summary, we describe distinct cellular and circuit mechanisms underlying processing of binocular visual information in mouse V1.

## Introduction

Sensory input to the cerebral cortex is processed in a hierarchical fashion (Mountcastle, 1978), with individual cortical neurons receiving input via multiple channels. Circuits and cells within the cortex need to isolate, integrate and route this information optimally for the brain to build up an accurate representation of the external world. Dissecting the circuit architecture and biophysical properties of the neurons involved is critical for understanding this process and will hopefully provide mechanistic insights into brain function.

The vertebrate cortex is divided into two hemispheres, with sensory input received and processed by the brain bilaterally. Early work in cats indicated that input from both eyes is processed by each hemisphere (Barlow et al., 1967; Hubel and Wiesel, 1962), yet we perceive a single view of the outside world (Welchman, 2016). Ultimately, this presumably requires that binocular visual input is combined at the level of single neurons in the brain. This process is not fully understood and is the focus of this study.

Visual information from the eyes reaches the visual cortex (V1) via the lateral geniculate nucleus (LGN). Retinal ganglion cell (RGC) axons from each eye project to the LGN in both hemispheres, with the amount of decussation at the optic chiasm dependent on the degree of binocular overlap in the visual field (Mason and Slavi, 2020). Work in mice indicates that LGN neurons are largely monocular, responding primarily to input from only one eye (Bauer et al., 2021). Binocularity in V1 is therefore thought to arise from convergent LGN input arriving in the cortex via separate, parallel pathways from the two eyes (Priebe and McGee, 2014). With this model, binocular information is processed separately and independently by V1 in each hemisphere.

In addition to this more traditional view, it has long been known that binocular V1 in the right and left hemisphere are reciprocally connected via callosal connections. In rodents, cats and primates these callosal connections are concentrated at the border between areas 17 and 18, a region of V1 responsible for processing binocular vision (Cusick and Lund, 1981; Kennedy et al., 1986; Olavarria, 2001). The importance of these callosal connections for binocularity in V1 is controversial, however. Some studies in cats, for example, have found that callosal projections are important for establishing ocular dominance in binocular neurons in V1 (Berlucchi and Rizzolatti, 1968; Blakemore et al., 1983; Yinon et al., 1992), whereas others indicate the opposite (Elberger and Smith, 1985; Minciacchi and Antonini, 1984). Similarly, some studies in rodents indicate that callosal projections carry ipsilateral eye input and play a significant role in shaping ocular dominance (Cerri et al., 2010; Dehmel and Lowel, 2014; Lee et al., 2019; Restani et al., 2009), whereas others have found evidence to the contrary (Coleman et al., 2009; Hagihara et al., 2021; Laing et al., 2015; Lewis and Olavarria, 1995; Olavarria and Van Sluyters, 1983; Olavarria, 2001; Ramachandra et al., 2020).

Much of the previous work investigating the functional role of callosal projections in rodents has relied on indirect measures of neuronal activity, such as extracellular field recording or calcium imaging, which may explain the different conclusions made by these studies. To address this issue, and to better understand the role of callosal projections in the processing of binocular visual information, here we combine *in vivo* and *in vitro* whole-cell patch-clamp electrophysiology, dual-colour optogenetics and 2-photon calcium imaging to identify the pathways and properties of neurons processing binocular visual information in mouse V1.

## Methods

### Animals

All experiments were approved by the Animal Experimentation Ethics Committee of the Australian National University in accordance with the Australian Code of Practice for the care and use of animals for scientific purposes. Wide type C57BL/6 mice of either sex as well as a limited number of transgenic mice expressing Cre in parvalbumin interneurons (PV-Cre mice) were used. Mice were housed in a controlled environment with a 12-hr light dark cycle with access to food and water ad libitum.

### Viral injections

Mice (4 to 6 weeks old) were placed in a chamber to induce light anaesthesia via brief exposure to isoflurane (3.5% in oxygen 1-2 L/min) then mounted in a stereotaxic frame with anaesthesia continued using isoflurane (1-1.5% in oxygen 1-2 L/min) delivered through a nose cone. A servo-controlled heating blanket (Harvard Instruments, USA) was used to maintain body temperature near 37°C. Ear bars were inserted into the ear canals to stabilize the head. The scalp was shaved with a hair-trimmer and any remaining hair removed with hair-removing cream. After cleaning the surgical area with 70% ethanol, an incision was made along the midline of the scalp using a single-edged scalpel blade to expose the skull. After cleaning and drying the exposed skull, lambda and bregma were marked as reference landmarks. A small craniotomy (0.5mm diameter) was then made using a dental drill over the injection site. Viral constructs were injected into layer 2/3 (L2/3) of binocular visual cortex (bV1; 0.5 mm anterior, 3.0 mm lateral to Lambda; depth: 0.2-0.4 mm) or the lateral geniculate nucleus (LGN; 2.15 mm posterior, 2.35 mm lateral to Bregma; depth: 2.7-2.9 mm) using borosilicate glass pipettes (Sutter), pulled on a microelectrode puller (Sutter Instruments, USA) and broken to a tip diameter around 10-15 μm. Injection pipettes were back-filled with mineral oil and front-loaded with viral suspension. The pipette containing the viral suspension was slowly lowered (minimising tissue damage) to the target depth and multiple boluses of virus (23 nl) were pressure injected using a Nanoject II (Drummond) at 5-minute intervals between injections. After the final injection, the pipette was left in place for an additional 5 min to allow the virus to diffuse before being slowly removed. The scalp was then sutured and Betadine applied to prevent infection of the wounded area. Mice were injected subcutaneously with 0.3 ml ketoprofen (1 mg/ml) for pain management and transferred to their home cage containing wet chow and water for recovery and monitored daily until the day of experiment. Mice were held for 3 to 6 weeks to allow viral expression prior to *in vivo* and *in vitro* experiments.

### Viral Vectors

For expression of ChR2 and ChrismonR a viral suspension containing AAV1-hSyn-ChR2(H134R)-eYFP-WPRE.hGH (0.1-0.4 µl) or AAV1-Syn-ChrimsonR-tdTomato (0.3-0.5 µl) was injected into V1 in one hemisphere. For optogenetic inactivation experiments, a viral suspension containing Cre-dependent ChR2 (AAV1-Ef1a-DIO-hChR2(E123A)-EYFP; 0.6-0.9 µl) was injected into V1 in one hemisphere of PV-Cre animals. For *in vivo* 2-photon calcium imaging, a viral suspension containing AAV1-CamKII-GCaMP6s-eYFP-WPRE.SV40 (0.2-0.3 µl) was injected into V1 in one hemisphere. For retrograde expression, a viral suspension containing AAVrg-hSyn-eGFP (0.2-0.4 µl) or AAVrg-hSyn-hChR2(H134R)-eYFP (0.4-0.5 µl) was injected into V1 in one hemisphere. For expression of ChrimsonR in contralateral V1 and ChR2 in the ipsilateral LGN AAV1-Syn-ChrimsonR-tdTomato (04-0.6 µl) was injected into V1 in one hemisphere and AAV1-hSyn-ChR2-EYFP (0.1-0.4 µl) was injected into the LGN in the opposite hemisphere. All data in the main results of the paper was obtained using AAVs sourced from Penn Vector Core (University of Pennsylvania). Some later control experiments described in the Supplementary data used AAVs sourced from Addgene (USA). While retrograde expression of AAV1 has been described previously (Zingg et al., 2017), this was not observed in experiments using AAVs from Penn Vector Core, consistent with earlier work from our laboratory (Gharaei et al., 2020). Retrograde expression of ChR2 and ChrismonR was observed, however, in later experiments using AAVs from Addgene, which was reduced to negligible levels following a 10-fold dilution with saline (Supplementary Figure 1).

### In vivo experiments

Mice (6 to 12 weeks old) were initially anesthetized by brief exposure to isoflurane (3.5% in oxygen 1-2L/min) with anesthesia subsequently maintained by intraperitoneal administration of urethane (0.5 to 1 g/kg) together with chlorprothixene (5 mg/kg). The level of anesthesia was regularly monitored by checking hind paw and corneal reflexes, and maintained at a stable level by administering top-up injections of urethane together with chlorprothixene as required (10% of original dose). Atropine (0.3 mg/kg, 10% w/v in saline) was administered subcutaneously to reduce secretions. The animal was placed on a servo-controlled heating blanket (Harvard Instruments, USA) to maintain a body temperature near 37°C. A custom-built head holder was glued to the skull and stabilized with dental cement. The head holder was mounted on a steel frame to minimize head movement. A craniotomy was performed above binocular V1 in one hemisphere (0.5 mm anterior, 3.0 mm lateral to Lambda). Whole-cell current-clamp recordings were obtained from L2/3 pyramdial neurons with a Multiclamp amplifier (Multiclamp 700A, Axon Instruments, USA) using methods described previously (Margrie et al., 2002). Patch pipettes were pulled from borosilicate glass and had open tip resistances of 5-7 MΩ when filled with an internal solution containing (in mM): 130 K-gluconate, 10 KCl, 10 HEPES, 4 MgATP, 0.3 Na2GTP, 15 Na2Phosphocreatine (pH 7.25 with KOH, osmolality ∼290 mOsm). Electrodes were inserted perpendicular to the craniotomy and lowered rapidly using a Sutter micromanipulator with high positive pressure (∼200 mmHg) to pass the dura mater. The pressure was then dropped to 30 mmHg and the pipette advanced at a speed of ∼2 µm/s while searching for neurons. Bridge balance and capacitance compensation were used to correct for the voltage drop across the series resistance during current injection and for filtering by the pipette capacitance. Hyperpolarizing and depolarizing current steps (−200 pA to +600 pA; intervals of 50 pA) were applied via the somatic recording pipette to characterize passive and active properties. The contralateral and ipsilateral eye were illuminated independently using brief (20 or 50 ms duration) full-field LED flashes (530 nm; ∼80 Lux, unless otherwise stated; 20∼100 interleaved trials) with custom-made LED “goggles” positioned a short distance (∼1.5cm) in front of the eyes. For *in vivo* optogenetic activation/inactivation another craniotomy was performed above binocular V1 in the opposite hemisphere on the ChR2-injected side and optogenetic activation of ChR2 performed using a high-power LED (470 nm, Thorlabs) coupled to a 200 μm diameter optic fiber placed just below the pia.

### In vitro experiments

Three to four weeks after viral injection, mice were deeply anesthetized with isoflurane (3% in oxygen) and immediately decapitated. The brain was quickly extracted and sectioned in a chilled cutting solution containing (in mM): 110 choline chloride, 11.6 N-ascorbate, 26 NaHCO3, 7 MgCl2, 3.1 Na-pyruvate, 2.5 KCl, 1.25 NaH2PO4, 0.5 CaCl2 and 10 glucose (pH = 7.4). Coronal slices at 300 µm thickness containing binocular V1 were prepared using a Leica Vibratome 1000S. Slices were incubated in an incubating solution containing (in mM): 92 NaCl, 2.5 KCl, 1.2 NaH2PO4, 30 NaHCO3, 20 HEPES, 3 Na-pyruvate, 2 CaCl2, 2 MgSO4 and 25 glucose at 35°C for 30 min, followed by incubation at room temperature for at least 30 minutes before recording. All solutions were continuously bubbled with carbogen (95% O2/5% CO2). Somatic whole-cell patch-clamp recordings were made under visual control from L2/3 pyramidal neurons in binocular V1 using an Olympus BX50 microscope with a 40 or 60x objective and infrared-differential interference contrast optics (Stuart et al., 1993). During recording, slices were constantly perfused at ∼2 ml/min with carbogen-bubbled artificial cerebral spinal fluid (ACSF) containing (in mM): 125 NaCl, 25 NaHCO3, 3 KCl, 1.25 NaH2PO4, 2 CaCl2, 1 MgCl2 and 25 glucose maintained at 30-34°C. Patch pipettes were pulled from borosilicate glass and had open tip resistances of 5-7 MΩ when filled with an internal solution containing (in mM): 130 K-gluconate, 10 KCl, 10 HEPES, 4 MgATP, 0.3 Na2GTP, 10 Na2phosphocreatine and 0.3% biocytin (pH 7.25 with KOH). All recordings were made in current-clamp using a BVC-700A amplifier (Dagan Instruments, USA). Bridge balance and capacitance compensation were used to correct for the voltage drop across the series resistance during current injection and for filtering by the pipette capacitance. Series resistance was below 15-20 MΩ. Hyperpolarizing and depolarizing current steps (−200 pA to +600 pA; intervals of 50 pA) were applied via the somatic recording pipette to characterize passive and active properties. All *in vitro* experiments were performed in the presence of gabazine (10 µM) to block inhibition mediated by GABAA receptors.

For photo-stimulation of ChR2-expressing neurons and axon terminals a 470 nm LED (Thorlabs) was mounted on the epi-fluorescent port of the microscope (Olympus BX50) providing wide-field illumination through the microscope objective. For some experiments, the red-shifted opsin ChrismonR was used for photo-stimulation instead of, or together with, ChR2. Photo-stimulation of ChrismonR was achieved by wide-field illumination via the epi-fluorescent port of the microscope using a 590 nm LED (Thorlabs). The timing, duration and strength of LED illumination were controlled by the data acquisition software. In general, for ChR2 axon terminal activation 470 nm LED duration was set to 2 ms and for ChrimsonR axon terminal activation 590 nm LED duration was set to 5 ms. Tetrodotoxin (TTX) (1 μM) and 4-aminopyridine (4-AP) (100 μM) were added to the external solution to isolate direct (monosynaptic) responses to LED stimulation. Application of 6,7-dinitroquinoxaline-2,3-dione (DNQX) (10 μM) and amino-5-phosphonovaleric acid (APV) (25 μM) was used in some experiments to isolate light-evoked responses due to synaptic release of glutamate from those due to activation of ChR2/ChrimsonR expressed in the recorded neuron.

### Two-photon axon imaging and visual stimulation

Two-photon calcium imaging of trans-callosal axons in binocular V1 was performed in L2/3 in the opposite hemisphere from that injected with GCaMP6 while presenting drifting gratings to the ipsilateral eye. That is, to the eye on the same side as two-photon imaging. The visual stimulus consisted of sinusoidal drifting gratings with a temporal frequency of 2 Hz and spatial frequency 0.05 cycles/degree (duration 1 second) and was presented to the ipsilateral eye using a gamma-corrected LCD screen (Longordo et al., 2013). Visual stimuli were generated in MATLAB using the Psychophysics Toolbox (Brainard, 1997) extension and custom written programs. Two-photon imaging was performed using a Thorlabs A-scope with a 16x objective (0.8 numerical aperture) and a Chameleon II laser (λ=920nm). ROIs corresponding to axons were detected using suite2p (Pachitariu et al., 2016) and subsequently combined based on their correlation coefficients.

### Histology

Neurons were morphologically reconstructed and analyzed using a biocytin staining protocol and neuron tracing software. Recorded neurons were filled with 0.5% (5 mg/ml) biocytin. Whole-cell recordings were maintained for at least 10-15 minutes to allow the biocytin to diffuse into the dendrites. After the end of recordings, pipettes were slowly removed to keep the soma intact. Immediately after completion of the recording, brain slices were fixed in 4% paraformaldehyde and then stored at 4°C overnight. The next morning, slices were washed three times with 0.1M phosphate buffered saline (PBS). For staining and avidin-horseradish peroxidase diffusion into cells, tissue slices were initially added to a PBS solution containing 1% hydrogen peroxide (H2O2; 10% absolute methanol+ 90% PBS) for 15 min to block endogenous peroxidases. Tissue slices were then added to a PBS solution containing 1% bovine serum albumin (BSA) and 0.3% Triton X-100 for 1 hour at room temperature to permeabilize the cell membrane and block non-specific binding. This solution was then removed and tissue slices were incubated in a solution containing the avidin-biotin peroxidase reaction solution (Vectastain Elite ABC Kit, Vector laboratories Inc., USA) for 24 hours at 4°C. Tissue slices were then washed 3 times with PBS and incubated in 0.05% DAB (10 mg/20 ml) in PBS + 0.01% H2O2 for 5-10 min for stain visualization. Individual tissues slices were then mounted on glass slides with Mowiol (polyvinyl alcohol, Merck) mounting solution. Mounted tissue slices were stored at 4°C and cells were visualized under a microscope (Zeiss; Axioskop II, USA). 3D computerized reconstructions were made with a Zeiss 63x/1.4 oil-immersion objective and Neurolucida software (v.7, Microbrightfield Inc., Williston, VT).

### Data acquisition and analysis

Voltage and current signals were filtered at 10 kHz using the amplifier built-in filter and digitized at a sample rate of either 20 or 50 kHz using an A-D converter (ITC-16 or ITC-18, Instrutech/HEKA, Germany) under the control of acquisition software (Axograph X, Axograph Scientific, Australia) running on an Apple Macintosh computer (iMAC-i7). Axograph was used for both data acquisition and primary analysis. Exported data were further analysed with custom scripts in Igor Pro8 (Wavemetrics, USA) and MATLAB. Input resistance (R) was calculated based on sub-threshold voltage responses to somatic current steps (−300 pA to +100 pA in 50 pA steps; 1 second duration), with the slope of the steady-state voltage (V) to current (I) relationship used to estimate R (R=ΔV/ΔI). For *in vivo* experiments, neurons were classified as receiving synaptic input from the ipsilateral or contralateral eye, or during optogenetic activation of the opposite V1, if the initial synaptic response (within the 100 ms after stimulus onset) had an average amplitude greater than 1 mV (typically 10 trials were averaged). For *in vitro* experiments, neurons were classified as receiving callosal input (“responders”) if the average synaptic response (10 trials) to photo-stimulation of callosal axons was greater that 1 mV at an LED intensity of 0.3 mW under control conditions or 1.4 mW (or greater) in the presence of TTX and 4-AP. EPSP ocular dominance index (ODI) was calculated based on contralateral and ipsilateral eye EPSP amplitude using (EPSPcontra - EPSPipsi) / (EPSPcontra + EPSPipsi). Neurons with an EPSP ODI less than 1 but greater than −1 were classified as binocular neurons, whereas neurons with an EPSP ODI of 1 or −1 were classified as monocular. The orientation selectivity index (OSI) was calculated as 1-circular variance (Mazurek et al., 2014; Ringach et al., 2002; Tan et al., 2011). A neuron that responds exclusively to a single orientation will have an OSI of 1, whereas neurons that respond equally to all orientations will have an OSI of 0. For paired data, Wilcoxon’s non-parametric signed-rank test, a paired t-test or a two-way analysis of variance (ANOVA) were used to test statistical significance. Statistical significance was set at p < 0.05. Results are presented as average values ± the standard error of the mean (SEM), unless otherwise stated. In the Figures “ns” denotes not statistically significant, whereas asterisks denote statistical significance (*P<0.05, **P<0.01, ***P<0.001, ****P<0.0001).

### Compartment modelling

Simulations were performed with the NEURON 7.4 simulation environment (Carnevale and Hines, 2006). A multicompartment model was obtained by reconstructing a biocytin-filled L2/3 pyramidal neuron (NeuroMorpho.org ID Martin, NMO_00904). The reconstructed neuronal model consisted of 112 compartments, subdivided into a total of 986 segments. We tuned the passive properties of the model to match ours and previous experimental findings (Larkum et al., 2007; Smith et al., 2013). The specific membrane resistance (Rm) and axial resistance (Ri) were tuned to match subthreshold voltage responses during somatic current injections with the input resistance of the model (90 MΩ) matched to that observed in experiments. The passive properties of the model were as follows: specific membrane resistance: Rm=9,000 Ω•cm^2^, specific membrane capacitance: Cm=1 μF/cm^2^ and internal resistance: Ri=150 Ω•cm. The resting membrane potential was set to −75 mV. The apparent membrane time constant was 9 ms. These values are similar to the average resting membrane potential and input resistance observed in experiments. Active conductances were included at axonal, somatic and dendritic compartments as follows (in mS/cm^2^): voltage-activated sodium channels (axon=3,000; soma=15; dendrites=3); delayed-rectifier (non-inactivating) potassium channels (axon=200; soma=20; dendrites=2); low-threshold voltage-activated calcium channels (soma=0.3e-3; dendrites=0.8e-3). Voltage-activated Na+ and K^+^ channels were included in the axon, soma and dendrites to match the experimentally observed somatic f/I relationship. Uniform densities of A-type potassium (Kv4.2) were used in the soma and dendrites (Bekkers, 2000; Korngreen and Sakmann, 2000). In addition, D-type (Kv1.1) channels were located in the axon and M-type (Kv7.2/3) channels were included at somato-dendritic locations at the indicated densities, consistent with experimental findings (Battefeld et al., 2014; Kole et al., 2007). High-frequency bursts of somatic action potentials can recruit dendritic electrogenesis in L2/3 cortical pyramidal neurons mediated by voltage-activated Ca^2+^ channels (Larkum et al., 2007). The frequency at which backpropagating action potentials lead to generation of dendritic Ca^2+^ spikes, known as the critical frequency (Larkum et al., 1999), is dependent on the dendritic Ca^2+^ channel density. The dendritic Ca^2+^ channel density was therefore tuned to match the critical frequency of 120 Hz observed experimentally in L2/3 neurons (Larkum et al., 2007).

## Results

### Distinct properties of ipsilateral and contralateral inputs to binocular V1

The vast majority (> 97%) of RGC axons from the rodent retina cross at the optic chiasm before reaching the LGN and V1 in the opposite hemisphere (Drager and Olsen, 1980; Lund, 1965). Therefore, we hypothesized that in rodents the majority of ipsilateral eye input to binocular V1 originates from callosal projections from the contralateral visual cortex rather than via the LGN on the same side, as suggested by earlier work (Cerri et al., 2010; Restani et al., 2009; Zhao et al., 2013). To investigate this, we made *in vivo* whole-cell current-clamp recordings from L2/3 pyramidal neurons in binocular V1 of anesthetised mice. EPSPs in L2/3 pyramidal neurons were evoked by brief (20 or 50 ms), full-field illumination of the ipsilateral or contralateral eye with LEDs (Fig. 1A,B). We found both binocular neurons, where EPSPs were evoked by stimulation of both the contralateral and the ipsilateral eye (94/130 neurons; 72%), as well as monocular neurons, where EPSPs were evoked by stimulation of one eye only (36/130; 27%; Fig. 1C). On average EPSPs evoked by stimulation of the contralateral eye were larger than those evoked by stimulation of the ipsilateral eye (Fig. 1D), with a range of EPSP ocular dominance indexes (ODIs) observed across the population of recorded neurons (Fig. 1E; n=130). These findings are consistent with earlier work were the ODI is based on spiking activity evoked by the contralateral and ipsilateral eye (Gordon and Stryker, 1996).

**Figure 1.**
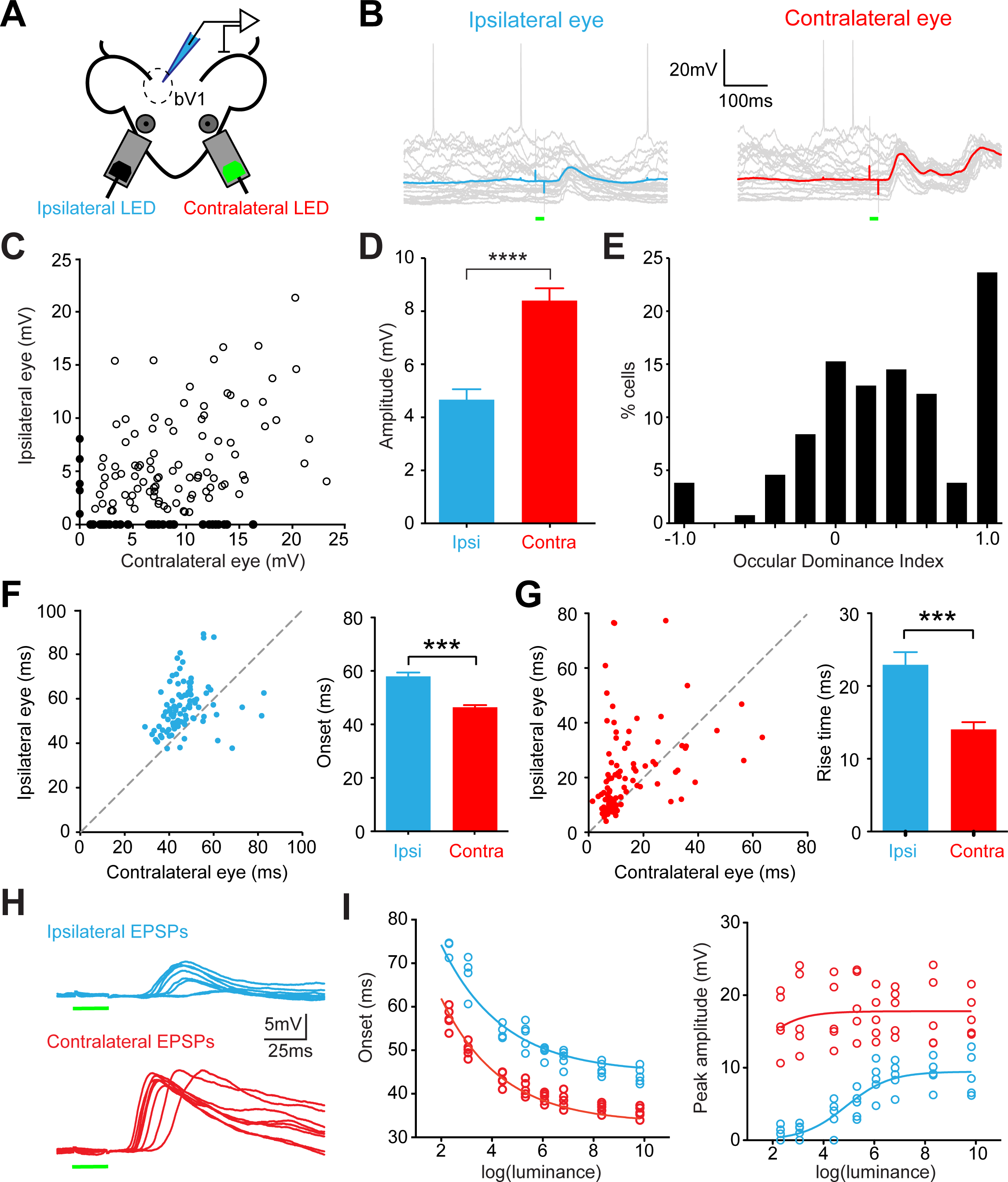
Distinct properties of ipsilateral and contralateral synaptic inputs to binocular neurons in visual cortex. **A)** Schematic of the experimental setup during *in vivo* whole-cell recording from binocular V1 (bV1) during contralateral or ipsilateral eye stimulation using LEDs positioned in front of the eyes to allow independent activation. The contralateral (green) LED is indicated as “on”. **B)** Synaptic responses of a binocular L2/3 pyramidal neuron to brief (20 ms) ipsilateral (left) and contralateral (right) LED eye stimulation (timing indicated by the green bar). Individual trials are shown in grey, averages in blue or red. **C)** Plot of contralateral eye versus ipsilateral eye EPSP amplitude in individual binocular (open symbols) and monocular (filled symbols) L2/3 pyramidal neurons in binocular V1 (n=130). **D)** Average contralateral and ipsilateral eye EPSP amplitude. **E)** Ocular Dominance Index of EPSP amplitude in L2/3 pyramidal neurons in bV1. **F)** Left, Plot of 20% onset time for contralateral versus ipsilateral eye EPSPs in different L2/3 pyramidal neurons. Right, Average (±SEM) 20% onset time of EPSPs for ipsilateral or contralateral eye stimulation (n=94 cells). **G)** Left, Plot of 20-80% rise time for contralateral versus ipsilateral eye EPSPs in different L2/3 pyramidal neurons. Right, Average (±SEM) 20-80% rise time of EPSPs for ipsilateral and contralateral eye stimulation (n=94 cells). **H)** EPSPs (average of 10 trials) in a binocular L2/3 pyramidal neuron during brief (20 ms) LED stimulation (green bars) of the ipsilateral (blue) or contralateral (red) eye at eight different LED intensities. LED artifacts blanked. **I)** Plots of EPSP onset (left; at 20% of peak) and EPSP peak amplitude (right) for EPSPs evoked by LED stimulation of the contralateral eye or the ipsilateral eye at different intensities.

Consistent with the hypothesis that ipsilateral eye input to binocular neurons arises via a different (longer) pathway compared to contralateral eye input, in the vast majority of binocular neurons (82/94; 87%) the onset time (at 20% peak amplitude) of the ipsilateral eye EPSP was significantly delayed relative to the contralateral eye EPSP (Fig. 1F; p<0:001; ipsi: 58 ± 1.4 ms vs. contra: 46 ± 0.8 ms; n=94). In addition, the rise time (20-80%) of the ipsilateral eye EPSP was significantly slower than the contralateral eye EPSP (Fig. 1G; p<0:001; ipsi: 23 ± 1.8 ms vs. contra:14 ± 1.0 ms; n=94). Ipsilateral eye EPSPs were delayed relative to contralateral eye EPSPs across all retinal illumination levels tested (Fig. 1H,I; average ipsilateral to contralateral onset time difference: 11.6 ± 0.6ms, p<0.05, non-linear regression). We also found that the dependence of ipsilateral and contralateral eye EPSP amplitude on the retinal illumination level was different (Fig. 1I, right).

These findings indicate that the properties of EPSPs evoked by brief activation of the ipsilateral eye are different from those generated by activation of the contralateral eye, with the response to activation of the ipsilateral eye delayed and slower compared to the contralateral eye. This suggests that inputs to binocular V1 from the ipsilateral and contralateral eyes are mediated via distinct pathways. This difference is unlikely to be due to differences in the synaptic location of ipsilateral and contralateral eye inputs on L2/3 neurons, as previous work indicates that ipsilateral and contralateral inputs co-localize onto the same dendritic branches of binocular L2/3 pyramidal neurons (Lee et al., 2019). Instead, the delay in the onset and slower rise time of synaptic responses evoked by activation of the ipsilateral eye is consistent with the idea that ipsilateral eye responses in binocular V1 arise primarily from callosal projections from the opposite V1, as this pathway would have a longer propagation delay compared to the conventional pathway via the LGN.

### Callosal projections from the opposite V1 convey ipsilateral eye information

Next, we asked whether the callosal input to binocular V1 from the opposite hemisphere carries ipsilateral eye information, as required if ipsilateral eye responses in binocular V1 arise from this callosal projection. To test this, we injected the genetically-encoded calcium indicator GCaMP6s (Chen et al., 2013) into V1 of one hemisphere and imaged callosal axons in the opposite hemisphere using two-photon calcium imaging while stimulating the ipsilateral eye with drifting gratings (Fig. 2A). Drifting gratings rather than LED flashes were used in these experiments to more robustly activate callosal axons. GCaMP6s-containing axons activated by visual stimulation of the ipsilateral eye were visualized in the binocular zone of the opposite V1 (Fig. 2B). Drifting gratings presented to the ipsilateral eye with different orientations evoked calcium responses in callosal axons that displayed strong orientation selectivity (Fig. 2C,D). Importantly, the vast majority of callosal axons (94%; 447/475) were responsive to ipsilateral eye stimulation (Fig. 2E) with a high orientation selectivity index (Fig. 2F). These data confirm that the callosal input to binocular V1 in the opposite hemisphere is conveying visual information from the ipsilateral eye.

**Figure 2.**
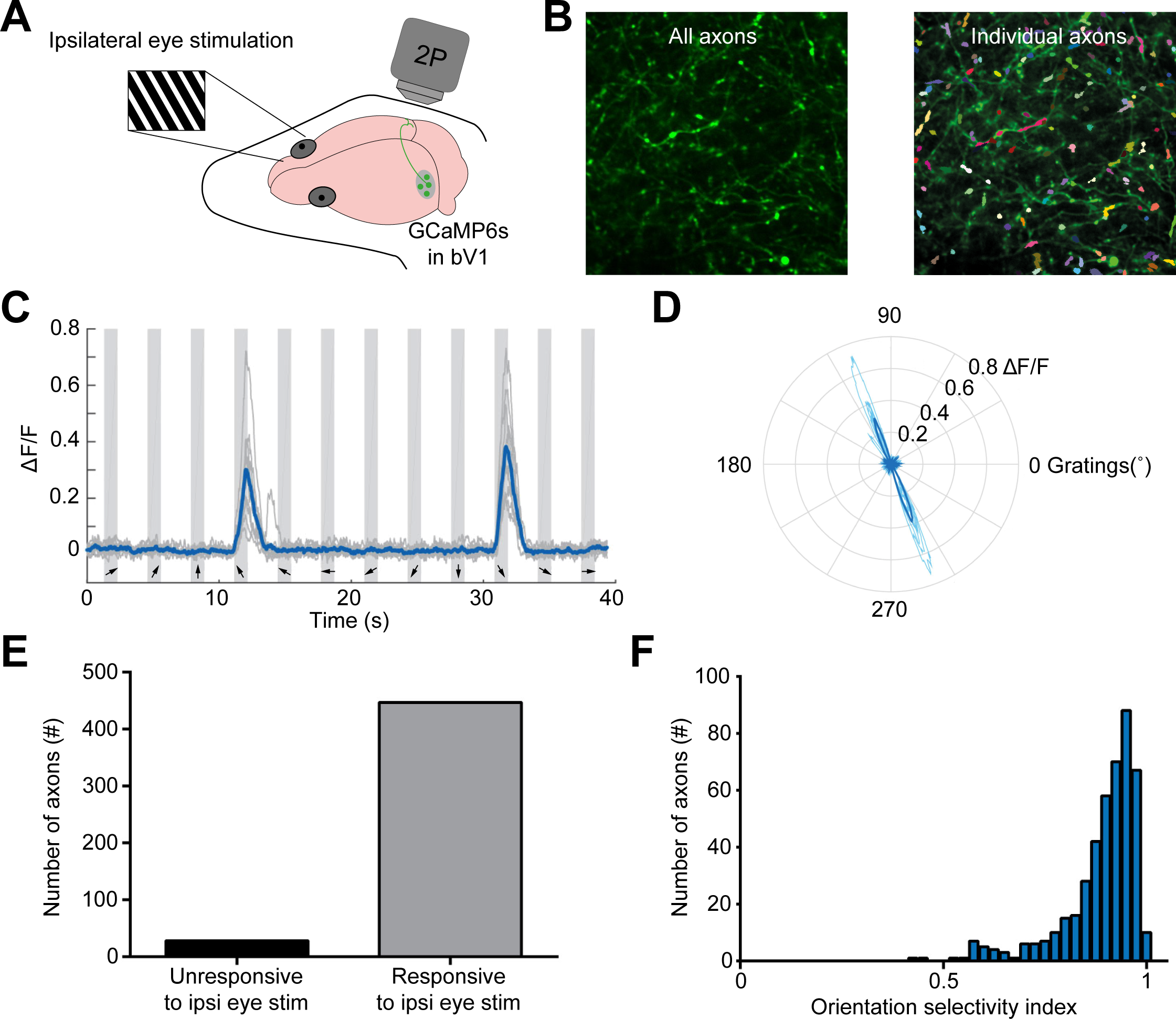
Callosal input from the opposite V1 to the binocular visual cortex carries ipsilateral eye information. **A)** Schematic of the experimental setup during GCaMP6s two photon (2P) calcium imaging of responses in callosal axons in the opposite binocular visual cortex during ipsilateral eye stimulation using drifting gratings. **B)** 2P image showing axons labeled with the calcium indicator (left) and sorted into individual color-coded axons with Suit2P software (right). **C)** Example of callosal axon changes in calcium florescence (ι1F/F) during presentation of ipsilateral drifting gratings at the indicated orientations (bottom arrows). **D)** Polar plot showing orientation tuning of the axon in **C**. **E)** Histogram of the proportion of axons responding to ipsilateral eye stimulation. **F)** Histogram of the orientation selectivity index for all recorded axons.

To investigate the importance of the callosal projection to ipsilateral eye responses in binocular neurons, we tested the impact of optogenetic inactivation of the callosal projection from the opposite V1 on ipsilateral eye responses L2/3 pyramidal neurons in binocular V1 (Fig. 3A). To inhibit callosal projections we injected Cre-dependent channelrhodopsin (ChR2) into V1 in one hemisphere of PV-Cre mice (Atallah et al., 2012). We then made *in vivo* whole-cell recordings from L2/3 pyramidal neurons in binocular V1 in the hemisphere opposite to the site of ChR2 injection and recorded the response to contralateral or ipsilateral eye stimulation with brief LED flashes with and without optogenetic activation of PV neurons in the opposite V1 (Fig. 3B). Control *in vitro* recordings in the ChR2-injected hemisphere indicated that optogenetic activation of fast-spiking interneurons in binocular V1 generated powerful inhibition of neighbouring L2/3 pyramidal neurons (data not shown). Optogenetic inhibition of the opposite V1 significantly reduced the amplitude of ipsilateral eye responses recorded *in vivo* in binocular neurons by approximately 50%, while having no impact on contralateral eye responses (Fig. 3C,D; p<0.001; n=18). These experiments provide functional evidence that a significant component (at least 50%) of the ipsilateral eye response in binocular V1 arises from the callosal projection from the opposite visual cortex.

**Figure 3.**
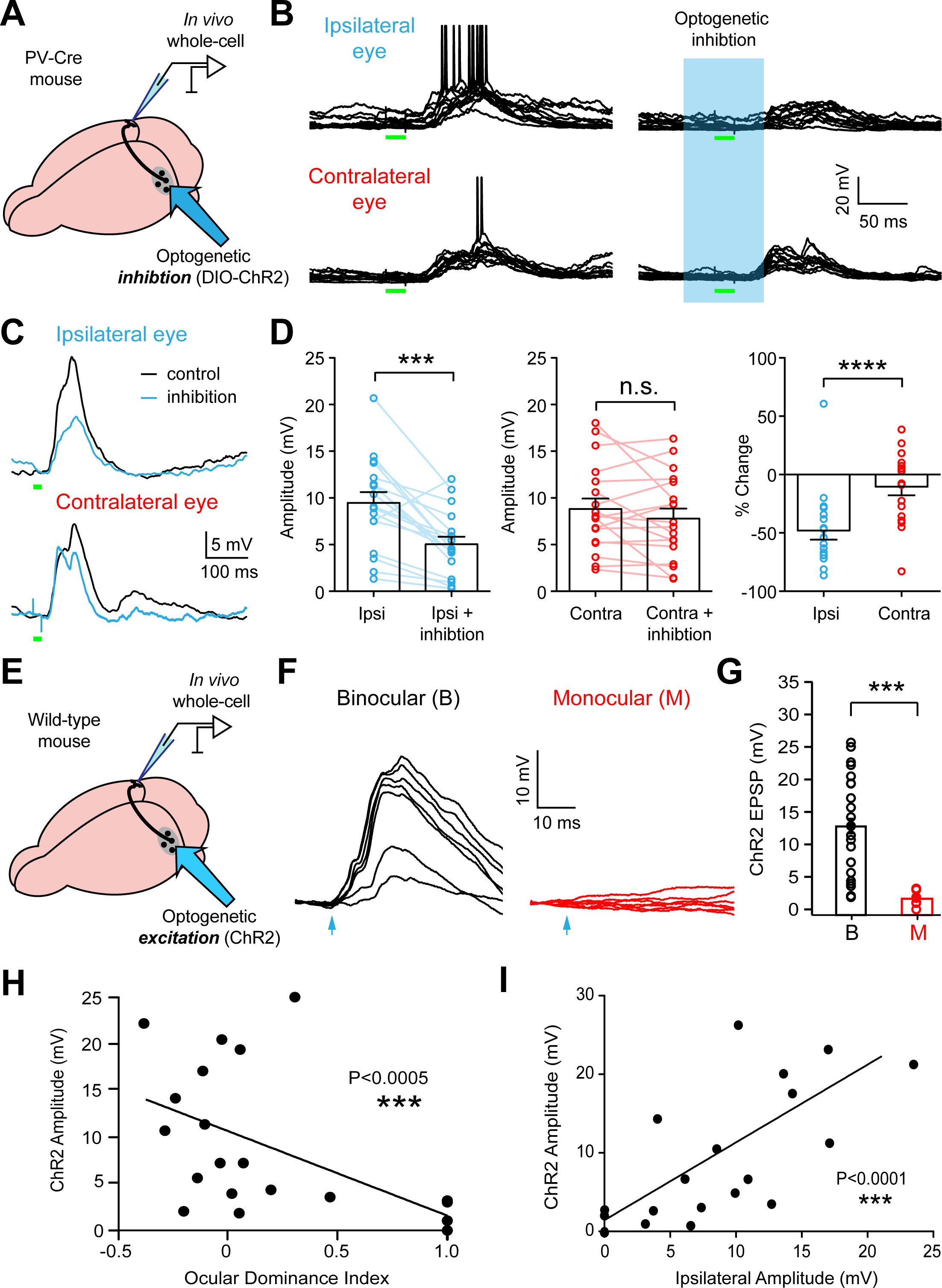
Impact of callosal input on responses in binocular and monocular neurons. **A)** Schematic of the recording situation during *in vivo* recording from binocular V1 combined with optogenetic inhibition of callosal input. **B)** Synaptic response of a binocular L2/3 binocular pyramidal neuron to brief (20 ms) ipsilateral (top) or contralateral (bottom) eye LED stimulation in control (left) and during optogenetic inhibition of the opposite hemisphere (cyan, right). **C)** Average synaptic response (20 trials) to ipsilateral (top) or contralateral (bottom) eye stimulation in control (black) and during optogenetic inhibition of callosal input (cyan). **D)** Individual and average amplitude (±SEM) of responses to brief (20 ms) ipsilateral (left) or contralateral (middle) eye LED stimulation during optogenetic inhibition of callosal input. Right, Individual and average (±SEM) percentage change in ipsilateral and contralateral eye LED responses following optogenetic inhibition of callosal input. **E)** Schematic of the recording situation during in vivo recording from binocular V1 combined with optogenetic excitation of callosal input. **F)** ChR2-evoked synaptic responses following brief (2 ms) optogenetic activation of callosal input (blue arrow) at different intensities (0 to 5.3 mW; 470 nm) in binocular (left) and monocular (right) neurons. **G)** Individual and average peak amplitude of ChR2 evoked responses in binocular (B; black) and monocular neurons (M; red; 5.3 mW). **H)** Plot of the peak amplitude of ChR2-evoked responses in individual neurons versus the synaptic ocular dominance index (ODI). Data fitted with a line. **I)** Plot of the peak amplitude of the ChR2-evoked synaptic response in individual neurons versus the amplitude of the ipsilateral eye response. Data fitted with a line.

### Only binocular neurons in V1 receive callosal input

We next determined whether binocular or monocular L2/3 pyramidal neurons (or both) receive callosal input. As the experiments above indicate that callosal input from the opposite V1 conveys ipsilateral eye input, we hypothesised that the neurons receiving callosal input will be binocular, whereas neurons that do not receive callosal input will be monocular. To test this idea, we injected ChR2 into V1 of one hemisphere and then performed *in vivo* whole-cell recordings from L2/3 neurons in binocular V1 in the opposite hemisphere (Fig. 3E). Optogenetic activation of the opposite V1 (470 nm; 2 ms; delivered via an optic fiber) was used to activate the callosal input. Functionally identified binocular neurons (n=24) were found to receive strong synaptic input from the contralateral hemisphere, whereas monocular neurons (n=8) did not (Fig. 3F). Even at the highest optogenetic intensities tested (up to 5.3 mW) monocular neurons did not show optogenetic responses to activation of the callosal projection from the opposite V1 (Fig. 3G; p<0.001; binocular: 12.5 *±* 0.6 mV vs. monocular: 0.98 *±* 0.3 mV). Importantly, we found that the amplitude of callosal synaptic responses was negatively correlated with the ODI, indicating that in individual neurons the strength of the callosal input is correlated with the degree of binocularity (Fig. 3H; slope=-9.1; p=0.0005). Furthermore, there was also a strong (almost one-to-one) correlation between the amplitude of the visual-evoked ipsilateral eye response and the callosal ChR2 response (Fig. 3I; slope=0.98; p<0.0001), indicating that cells with larger ipsilateral eye responses also received stronger callosal input. This correlation was not simply a consequence of differences in passive properties across the population, as it was preserved after both ipsilateral eye and ChR2 responses in individual neurons were normalized by the cell’s input resistance (Data not shown; slope=1; p<0.0001). Finally, we did not observe a correlation between the amplitude of the visual-evoked contralateral eye response and the callosal ChR2 response (Data not shown; slope=0.43; p=0.88). Together, these data indicate that binocular, but not monocular, neurons in binocular V1 receive callosal input from the opposite V1. Furthermore, the finding that the amplitude of ipsilateral, but not contralateral, eye responses is correlated with the strength of the callosal input further supports the idea that this pathway plays a critical role in conveying ipsilateral eye information to binocular neurons.

### Identification of monocular and binocular neurons in vitro

As only binocular neurons in binocular V1 receive callosal input from the opposite V1, we used the presence or absence of callosal input to identify binocular and monocular neurons *in vitro* and characterise their properties. To do this, we expressed ChR2 in V1 in one hemisphere and then performed whole-cell current-clamp recordings from L2/3 pyramidal neurons in coronal brain slices of the opposite binocular V1. Consistent with earlier studies (Cusick and Lund, 1981; Jacobson, 1970; Mizuno et al., 2007; Olavarria and Van Sluyters, 1983), we found that the axons from neurons in V1 in one hemisphere projected to the binocular region of V1 in the opposite hemisphere, i.e. to a region between the boundary of monocular V1 and secondary visual cortex (V2), which we call the “binocular band” (Fig. 4A). Recordings from L2/3 pyramidal neurons in V1 in the ChR2-injected hemisphere confirmed that 470 nm light evoked spiking (Fig. 4B, left), whereas low-power LED activation (470 nm, 2 ms, 0.3 mW) evoked putative synaptic responses in L2/3 pyramidal neurons in the binocular band in the opposite hemisphere (Fig. 4B; right). Putative synaptic responses were blocked by bath application of DNQX plus APV to block AMPA and NMDA receptors, confirming that they were synaptic in origin and not due to retrograde expression of ChR2 (Fig. 4C).

**Figure 4.**
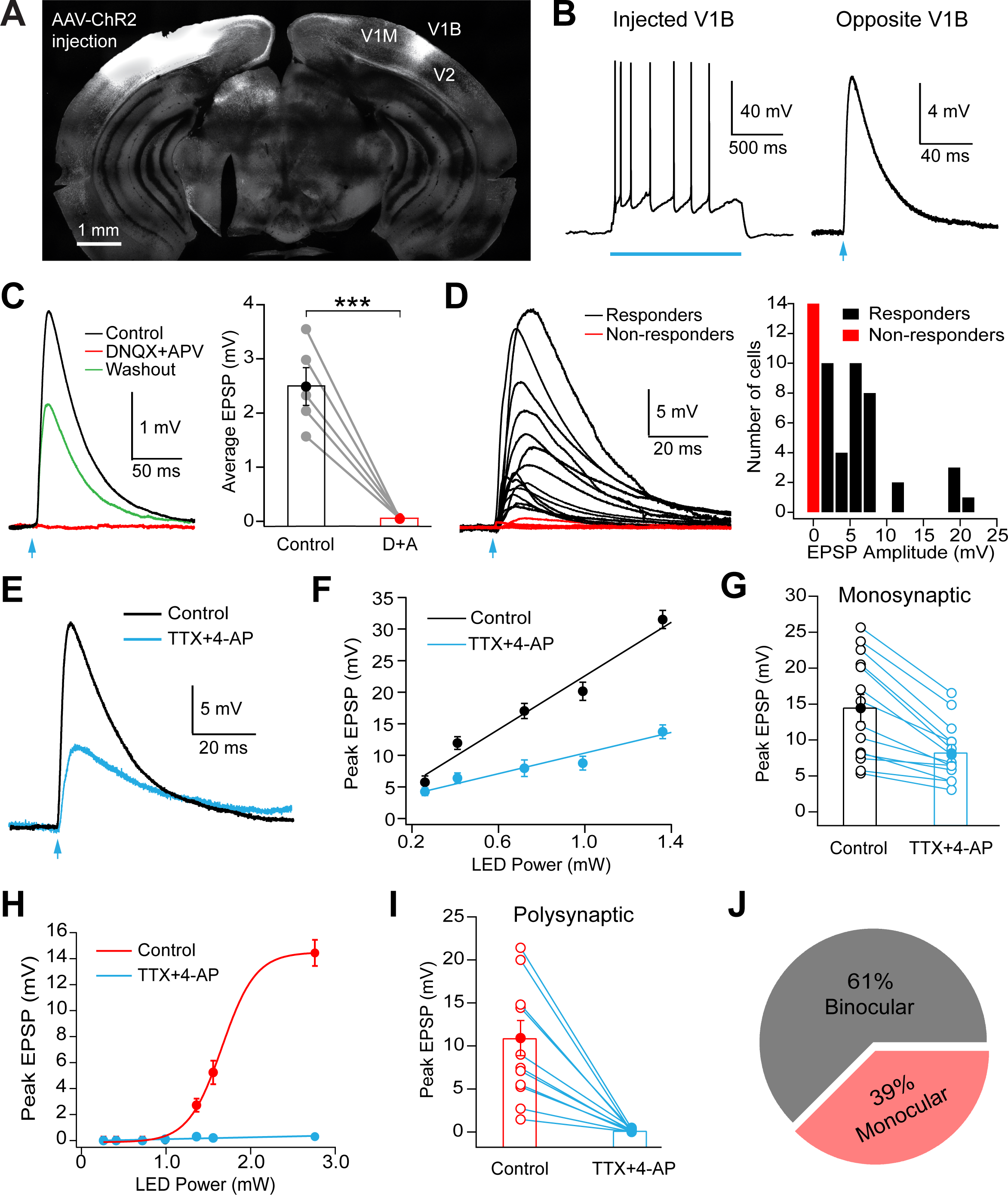
Different populations of L2/3 pyramidal neurons receive callosal input from the opposite V1. **A)** Confocal image of a coronal section showing expression of ChR2-eYFP in the injected V1 hemisphere (left) and long-range callosal axons on the opposite V1 (right) targeting the binocular zone at the border of V1 and V2. **B)** Left, Light-evoked action potentials in a L2/3 pyramidal neuron in the left (injected) hemisphere (0.3 mW; 1 s; blue bar). Right, Light-evoked EPSP in a L2/3 pyramidal neuron in the binocular zone of the right hemisphere to activation of callosal input from the opposite V1 (0.3 mW, 2 ms; blue arrow). **C)** Left, Light evoked EPSP in the binocular zone in control (black), DNQX+APV (red) and washout (green). Right, Pooled data showing the impact of DNQX+APV on EPSP amplitude (0.3 mW, 2 ms). **D)** Left, Light-evoked EPSPs in different L2/3 pyramidal neurons in the binocular zone (0.3 mW, 2 ms). Neurons with synaptic responses were classified as “responders” (black), whereas neurons that did not have a synaptic response (response < 1 mV) were classified as “non-responders” (red). Right, Histogram of callosal synaptic response amplitude for the responder and non-responder population (0.3 mW, 2 ms). **E)** Light-evoked (2.72 mW, 2 ms) synaptic response in control (black) and TTX+4-AP (blue; 2.72 mW, 2 ms) in a neuron receiving monosynaptic input. **F)** Synaptic response amplitude as a function of LED power in control and TTX+4-AP for the neuron shown in (**E**). **G)** Pooled data for EPSP peak amplitude in control (black) and TTX+4-AP (blue) in neurons receiving monosynaptic input (1 mW, 2 ms). **H)** Synaptic response amplitude as a function of LED power in control and TTX+4-AP for the neuron receiving polysynaptic input. **I)** Pooled data for EPSP peak amplitude in control (black) and TTX+4-AP (cyan) in neurons receiving polysynaptic input (2.72 mW, 2 ms). **J)** Pie chart showing the proportion of binocular and monocular neurons in binocular V1 based on the presence or absence of callosal input, respectively.

Based on the amplitude of synaptic responses, we classified L2/3 pyramidal neurons as either “responders” if EPSPs were greater than 1 mV or “non-responders” if EPSPs were less than 1 mV (Fig. 4D) for optogenetic LED activation at 0.3 mW (2 ms). To ensure that the absence of synaptic input was not due to poor ChR2 expression, we only included data from brain slices where at least one recorded neuron had a synaptic response to optogenetic activation at the lowest LED power tested (0.16 mW; 2 ms duration). In addition, we always checked that slices had a visible fluorescence band in the region between V1 and V2, indicating the presence of ChR2-expressing axons, and only recorded from L2/3 neurons within this region of the slice.

To test whether synaptic responses in the “responder” population were due to a direct, monosynaptic projection from the opposite V1, we bath applied TTX plus 4-AP (Petreanu et al., 2009). TTX blocks action potential initiation, abolishing synaptic responses due to activation of surrounding neurons, whereas 4-AP enhances transmitter release. Under these recording conditions synaptic responses can only be evoked by direct (monosynaptic) callosal input. Synaptic responses in the majority (81%) of “responders” remained in the presence of TTX plus 4-AP, indicating that this population of L2/3 pyramidal neurons receives monosynaptic callosal input from the opposite V1 (Fig. 4E-G). Using high-power LED stimulation (2.7 mW) synaptic responses could be evoked in the “non-responder” population, but these responses were blocked in the presence of TTX plus 4-AP, indicating that they were due to polysynaptic input (Fig. 4H,I). Overall, approximately 60% of L2/3 pyramidal neurons were found to receive monosynaptic callosal input from the opposite V1 and were classified as binocular neurons, whereas approximately 40% did not and were classified as monocular neurons (Fig. 4J).

### Properties of binocular and monocular neurons

We next characterized the passive, active and morphological properties of binocular and monocular L2/3 pyramidal neurons *in vitro*. In all cases the presence or absence of monosynaptic callosal input was confirmed in the presence of TTX plus 4-AP to determine if neurons were binocular or monocular. To investigate whether there were differences in passive properties we injected current steps and recorded subthreshold steady-state voltage responses to measure the input resistance of monocular and binocular neurons. Under control conditions (in the absence of TTX plus 4-AP) we found no significant difference in the input resistance or resting membrane potential between binocular and monocular neurons (Supplementary Fig. 2; n=59). In addition, morphological reconstruction of biocytin-filled binocular and monocular neurons showed no significant differences in their dendritic branching pattern (Supplementary Fig. 3; n=16). These data suggest that the passive electrical properties as well as the morphology of binocular and monocular neurons are similar. These observations also verify that the absence of callosal input was not simply due to a higher leak conductance in monocular neurons or a result of their dendritic tree being cut off during slicing.

Next, we focused on the active properties of binocular and monocular L2/3 pyramidal neurons. We first recorded active neuronal properties under control conditions and then perfused the slice with TTX plus 4-AP to determine if the recorded neuron received direct, monosynaptic callosal input or not. As with the above experiments, the presence or absence of direct, monosynaptic callosal input in TTX plus 4-AP was used to classify neurons as binocular or monocular, respectively. With regard to single action potential properties, there was a trend for increased action potential half-widths in binocular compared to monocular neurons this was not significant (Fig. 5A; n=35). Consistent with this trend, the after-hyperpolarizing potential (AHP) in binocular neurons was found to be smaller than that in monocular neurons (Fig. 5B; n=35). These data indicate differences in the conductances associated with action potential repolarization between binocular and monocular neurons. Next, we investigated the dependence of action potential firing on the magnitude of somatic current injection in binocular and monocular neurons: the so-called input/output function or f/I curve. This analysis indicated a significant difference between the f/I curve in binocular and monocular neurons (Fig. 5C). We extracted three parameters from the f/I curve, namely: action potential firing rate for a 600 pA current step, slope of the f/I curve and the f/I curve midpoint. While we observed no significant difference in the f/I curve midpoint, action potential firing rate and f/I slope were both significantly lower in binocular compared to monocular neurons (Fig. 5D-F; p<0.001; n=35). Furthermore, we found a significant negative correlation between both action potential firing rate (at 600 pA) and f/I slope recorded under control conditions with the amplitude of the callosal EPSP in TTX plus 4-AP (Fig. 5G,H; p<0.001). Similar results were obtained in mice injected with 1:10 diluted AAV1-ChR2 (Supplementary Fig. 4). These data suggest that binocular neurons are less excitable than monocular neurons and that the decreased excitability in binocular neurons is correlated with the strength of the callosal input. Consistent with this idea, the excitability of functionally identified monocular and binocular neurons *in vivo* was different, with the rate of action potential firing to a 600 pA current step negatively correlated with ipsilateral, but not contralateral, eye EPSP amplitude (Supplementary Fig. 5). These *in vivo* experiments validate our observations *in vitro*, and together indicate that binocular neurons have reduced excitability compared to monocular neurons, with the reduction in excitability correlated with the amplitude of ipsilateral eye input.

**Figure 5.**
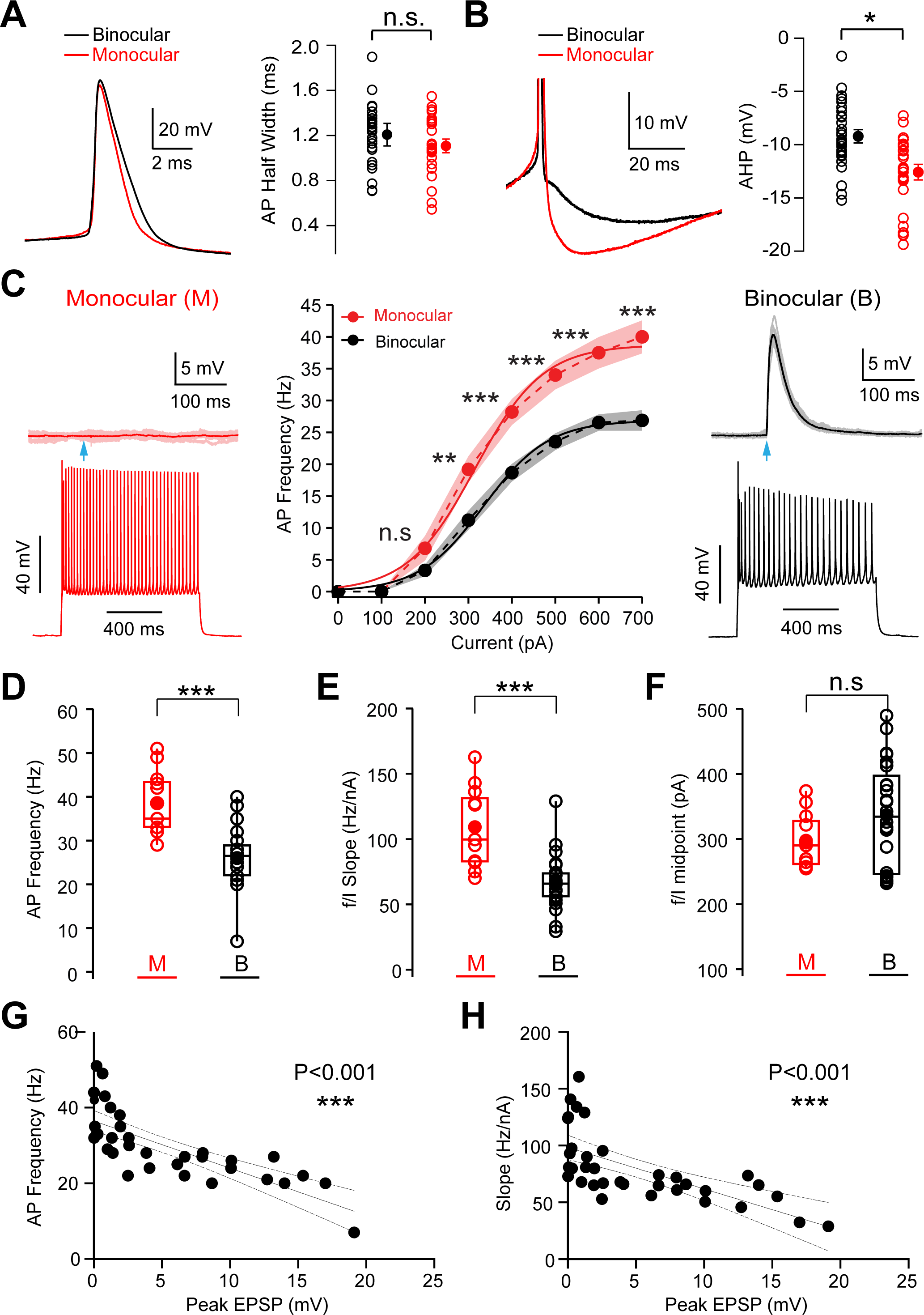
Differences in the active properties of binocular and monocular neurons. **A)** Left, Example of an action potential (AP) in a binocular (black) and monocular (red) L2/3 pyramidal neuron under control conditions. Right, Pooled data showing AP half width for all binocular (black) and monocular neurons in control (red). **B)** Left, Example of the after-hyperpolarization potential (AHP) following an AP in a binocular (black) and monocular (red) L2/3 pyramidal neuron under control conditions. Right, Pooled data showing AHP amplitude (relative to AP threshold) for all binocular (black) and monocular neurons in control (red). **C)** Left & Right, Example of the synaptic response to callosal input in TTX plus 4-AP (top) and AP firing in response to a 600 pA current pulse in control (bottom) in monocular (left, red) and binocular (right, black) L2/3 pyramidal neurons. Middle, Average AP firing frequency versus current injection amplitude (f/I) curve in binocular (black) and monocular (red) neurons under control conditions. Data fit with sigmoidals. Shading indicates SEM. **D-F)** Pooled data showing the AP firing frequency (**D**; 600 pA current pulse), f/I slope (**E**) and f/I midpoint (**F**) in binocular (black) and monocular (red) L2/3 pyramidal neurons under control conditions. Open circles indicate values from each cell and closed circles indicate mean. **G,H**) Relationship between the amplitude of the callosal input in the presence of TTX plus 4-AP and AP firing frequency (**G**; 600 pA current pulse) and sigmoid fit slope (**H**) recorded under control conditions.

### Binocular neurons express D-type potassium channels

To understand the biophysical mechanisms underlying the excitability difference between binocular and monocular neurons, we tuned the active and passive properties of a morphologically realistic L2/3 neuron model to match our experimental observations (Supplementary Fig. 6). Previous work indicates that slowly-activating and non-inactivating K^+^ conductances play an important role in regulating neuronal excitability (Bekkers and Delaney, 2001; Metz et al., 2007; Ordemann et al., 2019). Kv1.1/2 channels are localized at high densities in the axon initial segment of cortical neurons (Kole et al., 2007; Lorincz and Nusser, 2008). Therefore, we investigated how changes in the density of axonal Kv1.1 channels affect the f/I curve, with an aim to mimic the experimental difference in f/I curves between binocular and monocular neurons. These simulations revealed that the f/I curve in the model was highly sensitive to the density of axonal Kv1.1/2 channel (Supplementary Fig. 6J). The f/I curve in the model was also sensitive to the density of somato-dendritic Kv7.2/3 channels (Supplementary Fig. 6K). The influence of changes in the density of Kv1.1/2 and Kv7.2/3 channels on the maximum action potential firing frequency, f/I slope and f/I midpoint were approximately linear. The slope of these linear fits gives a measure of how sensitive the f/I curve is to a change in Kv1.1/2 and Kv7.2/3 channel density. This analysis indicated that much larger changes in Kv7.2/3 channel density compared to Kv1.1/2 channel density were required to have the same impact on f/I curve parameters, whereas changes in the dendritic and somatic A-type K+ channel density did not have a significant impact on f/I curve parameters (data not shown). In summary, our model predicts that the biophysical mechanism underlying the difference in intrinsic excitability between binocular and monocular neurons is most likely to be due to a difference in Kv1.1/2 channel density.

To test whether cell-specific differences in Kv1.1/2 (D-type) K^+^ channel expression underlie the observed difference in intrinsic excitability between binocular and monocular L2/3 pyramidal neurons, we investigated the impact of blocking these channels with a low concentration (300 µM) of 4-AP (Bekkers and Delaney, 2001). Neurons in these experiments were putatively identified as binocular or monocular based on the presence or absence, respectively, of synaptic responses to callosal input during optogenetic activation at low intensities (0.3 mW LED power) under control conditions. These experiments indicated that bath application of low concentrations of 4-AP led to a significant increase in action potential firing in binocular neurons (Fig. 6A), whereas it had little impact on monocular neurons (Fig. 6B). Quantification of these experiments indicated that low concentrations of 4-AP led to a significant increase in the maximum firing rate and f/I slope of binocular neurons by 28% and 35%, respectively, but no significant change in maximum firing rate or f/I slope of monocular neurons (Fig. 6C,D). Importantly, following application of 4-AP (300 µM) the maximum firing rate and f/I slope of binocular neurons was not statistically different from monocular neurons in control (Fig. 6C,D). Similar observations were made using α-dendrotoxin (DTX), a specific blocker of Kv1.1/2 channels (Harvey, 2001; Robertson et al., 1996; Stuhmer et al., 1989). Bath application of DTX (500 nM) increased the maximum firing rate and f/I slope of binocular neurons, while having no significant impact on monocular neurons (Supplementary Fig. 7). Importantly, as with application of low concentrations of 4-AP, application of DTX abolished the difference in maximum firing rate and f/I slope between monocular and binocular neurons (Supplementary Fig. 7). Together, these data support the idea that differences in the excitability of binocular and monocular neurons are due to differences in the expression of Kv1.1/2 channels, with the expression of Kv1.1/2 channels substantially higher in binocular neurons compared to monocular neurons, leading to reduced excitability.

**Figure 6.**
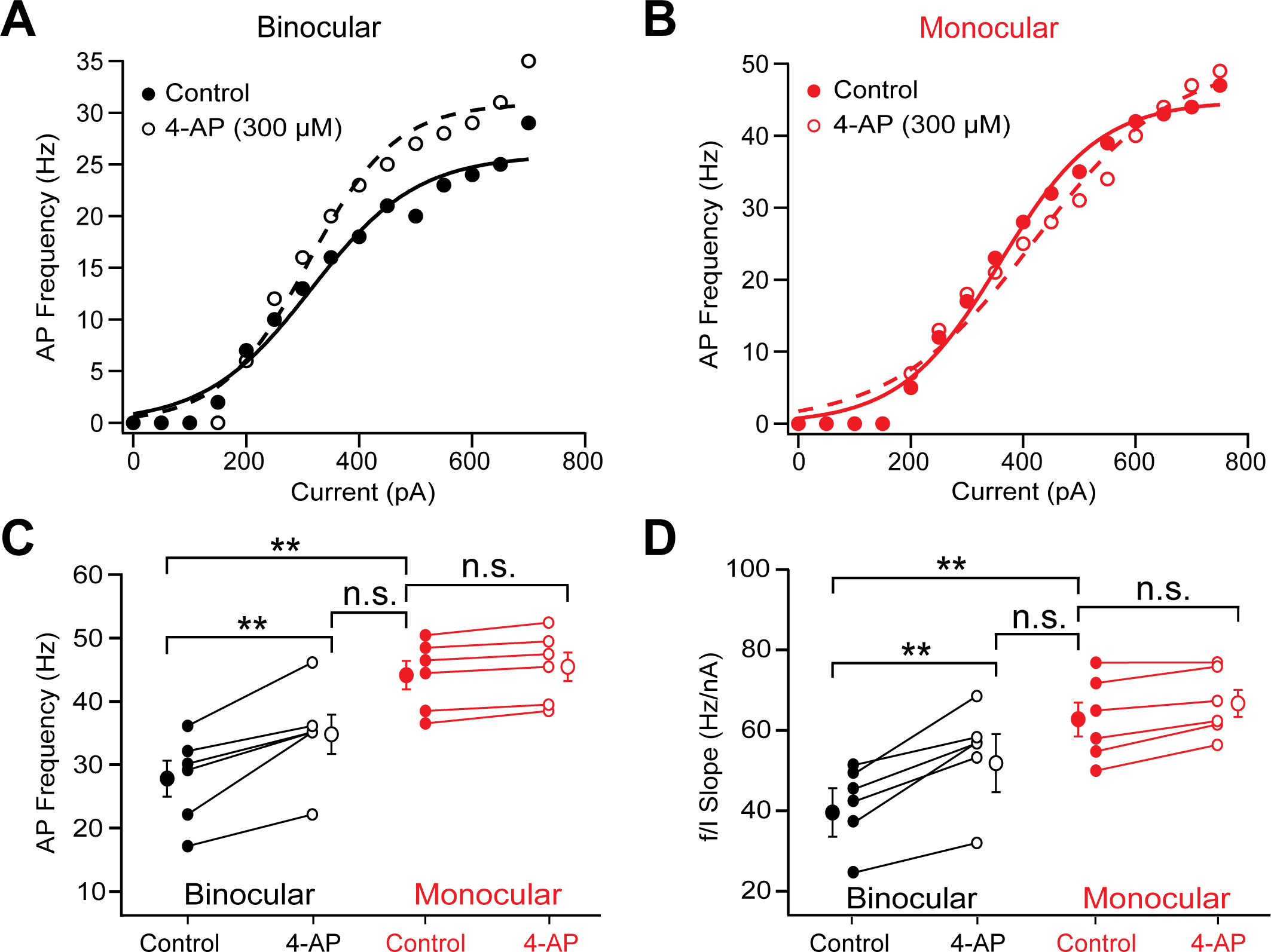
Reduced excitability in binocular neurons is due to higher expression of Kv1 channels. **A,B)** Action potential (AP) firing frequency versus current amplitude (f/I) curves in control (closed circles) and after application of a low concentration of 4-AP (open circles) in putative binocular neurons receiving callosal input (**A**) and monocular neurons which did not (**B**). **C,D)** Individual and average data for AP firing frequency during 600 pA current pulses (**C**) and f/I slope (**D**) obtained from sigmoid fits to data like that shown in (**A**,**B**) in control and after application of a low concentration of 4-AP in binocular (black) and monocular (red) neurons.

### Monocular neurons send callosal projections

We next determined whether binocular or monocular neurons in binocular V1 (or both) send callosal projections to the opposite V1. To investigate this, we co-injected a retrograde AAV (Tervo et al., 2016) tagged with eGFP and the long wavelength ChR2 variant ChrimsonR into V1 in one hemisphere (Fig. 7A). We then made whole-cell current-clamp recordings from L2/3 pyramidal neurons in the opposite V1 in the presence of TTX plus 4-AP (Fig. 7A). Both retrogradely labeled (eGFP-positive) and non-retrogradely labeled (eGFP-negative) L2/3 pyramidal neurons were visually identified within the binocular zone of V1 (Fig. 7B). Importantly, we found that following optogenetic activation of callosal input (590 nm; 5 ms) only eGFP-negative neurons received callosal input, whereas eGFP-positive neurons did not (Fig. 7C-E). As only binocular neurons receive callosal input from the opposite hemisphere, these data suggest that monocular, but not binocular, neurons send a callosal projection to the opposite V1. Importantly, the observation that retrogradely labelled L2/3 pyramidal neurons did not respond to ChrismonR activation indicates that AAV1-ChrismonR was not retrogradely transported in these experiments.

**Figure 7.**
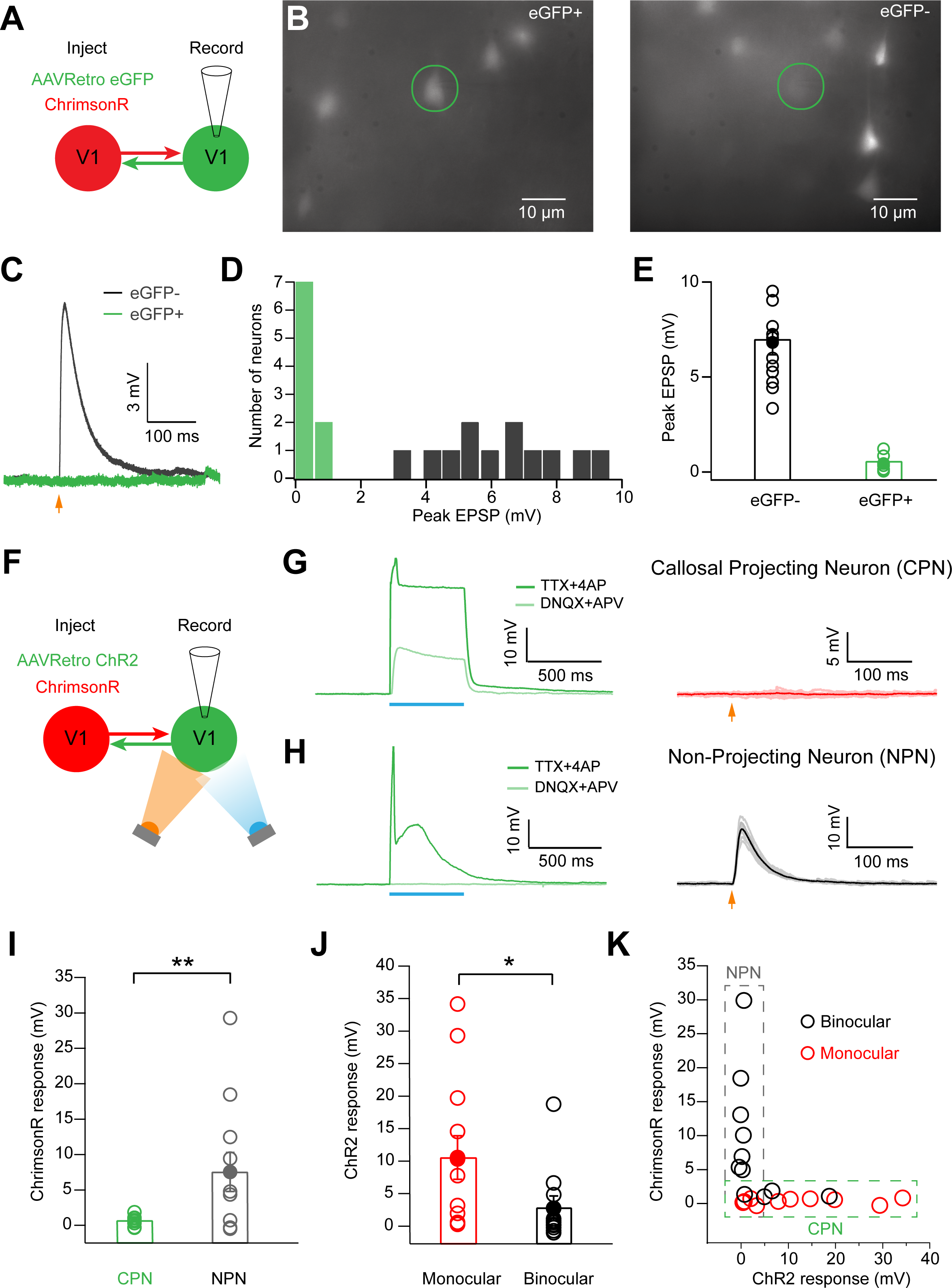
Only monocular neurons send callosal projections to the opposite V1. **A)** Schematic of the experimental setup showing ChrimsonR and AAVretro-eGFP injection into V1 in one hemisphere and recording in the opposite V1. **B)** Fluorescent image of a retrogradely labelled eGFP positive (eGFP+; left; green circle) and eGFP negative (eGFP-; right; green circle) L2/3 pyramidal neuron in binocular V1. **C)** Synaptic responses in TTX and 4-AP following activation of callosal input (orange arrow) in eGFP+ (green) and eGFP-(black) neurons. **D)** Histogram of callosal EPSP peak amplitude in TTX plus 4-AP in retrograde positive (eGFP+; green) and negative (eGFP-; black) L2/3 pyramidal neurons in binocular V1. **E)** Summary of callosal EPSP peak amplitude in TTX plus 4-AP in retrograde positive (eGFP+; green) and negative (eGFP-; black) L2/3 pyramidal neurons in binocular V1. **F)** Schematic of the experimental setup showing ChrimsonR and AAVretro-ChR2 injection into V1 in one hemisphere and recording and optical stimulation in the opposite V1. **G)** Left: Voltage response of a callosal projecting L2/3 pyramidal neuron (CPN) to a prolonged (500 ms) 470 nm light pulse (blue bar; 5 mW) in the presence of TTX plus 4-AP and after wash in of DNQX plus APV. Right: Synaptic response of the same neuron to activation of callosal axons by a brief (5 ms) 590 nm light pulse (2.8 mW) in the presence of TTX plus 4-AP to activate ChrimsonR (orange arrow). **H)** Left: Voltage response of a non-callosal projecting L2/3 pyramidal neuron (NPN) to a prolonged (500 ms) 470 nm light pulse (blue bar; 5 mW) in the presence of TTX plus 4-AP and after wash in of DNQX plus APV. Right: Synaptic response of the same neuron to activation of callosal axons by a brief (5 ms) 590 nm light pulse (2.8 mW) in the presence of TTX plus 4-AP to activate ChrimsonR (orange arrow). **I)** Pooled data of the peak amplitude of ChrimsonR-evoked synaptic responses in the CPN and NPN population, defined by the presence or absence, respectively, of a ChR2 response in DNQX plus APV. **J)** Pooled data of the amplitude of ChR2-evoked responses in the presence of DNQX plus APV in binocular (black) and monocular (red) neurons, defined by the presence or absence, respectively, of ChrimsonR responses indicating callosal input. **K)** Peak amplitude of ChrimsonR callosal input in the presence of TTX plus 4-AP plotted against the peak amplitude of ChR2-evoked responses in the presence of DNQX plus APV for binocular (black) and monocular (red) L2/3 pyramidal neurons in binocular V1.

One caveat with this experiment is that optical ambiguity in the identification of retrogradely positive and negative cells could lead to errors in visually classifying neurons as eGFP-positive or negative. To overcome this issue, we co-injected retrograde ChR2 and ChrimsonR into V1 in one hemisphere and again made whole-cell current-clamp recordings from L2/3 pyramidal neurons in the binocular band of the opposite hemisphere (Fig. 7F). The advantage of this approach is that it allowed us to electrically tag callosal projecting neurons based on the presence of retrograde ChR2 expression, eliminating any potential optical bias. In the presence of TTX plus 4-AP we then recorded the response of L2/3 pyramidal neurons to a prolonged blue light pulse (470 nm; 500 ms), to activate retrograde ChR2, and a brief orange light pulse (590 nm; 5 ms) to activate ChrimsonR in callosal axons. We then washed in DNQX (an AMPA receptor blocker) and APV (an NMDA receptor blocker) to block glutamatergic synaptic responses to determine which neurons directly express ChR2.

Using this protocol, we identified two populations of neurons: one population where the response to blue light-activation remained in the presence of DNQX and APV (Fig. 7G, left), indicating retrograde ChR2 expression, and another where the response was abolished, indicating an absence of direct ChR2 expression (Fig. 7H, left). These two populations represent callosal projecting and non-projecting neurons, respectively. Importantly, optogenetic activation of ChrimsonR did not evoke synaptic responses in the callosal projecting population (Fig. 7G, right), whereas synaptic responses were observed in the non-projecting population (Fig. 7H, right). We next quantified the peak amplitude of ChrimsonR synaptic responses in the presence of TTX plus 4-AP, as well as the amplitude of the retrograde ChR2 response in the presence of DNQX plus APV. Callosal projection neurons (CPNs) that had a retrograde ChR2 response in the presence of DNQX plus APV had significantly smaller ChrimsonR synaptic responses than non-projection neurons (NPNs) that did not have a retrograde ChR2 response in the presence of DNQX plus APV (Fig. 7I). In addition, the retrograde ChR2 response in the presence of DNQX plus APV was significantly larger in monocular neurons that did not receive callosal input compared to binocular neurons that did (Fig. 7J). We then plotted the amplitude of the retrograde ChR2 response in the presence of DNQX plus APV against the amplitude of the ChrimsonR synaptic response in the presence of TTX plus 4-AP for each cell individually (Fig. 7K). This revealed essentially non-overlapping distributions of callosal projecting, monocular neurons and non-projecting, binocular neurons, consistent with the idea that these two populations are distinct. Together, these data are consistent with our findings using fluorescent retrograde labelling to identify callosal projecting neurons, and provide further support for the idea that monocular, but not binocular, neurons send a callosal projection to the opposite V1.

### Convergence of thalamic and callosal input in binocular V1

To investigate convergence of ipsilateral and contralateral eye inputs onto L2/3 pyramidal cells in binocular V1 we applied a dual-opsin approach (Hooks et al., 2015), expressing ChrimsonR in the opposite V1 and ChR2 in the ipsilateral LGN. Significant overlap of LGN axons (expressing eYFP) and callosal axons (expressing tdTomato) was observed in the binocular zone of V1 (Fig. 8A).

**Figure 8.**
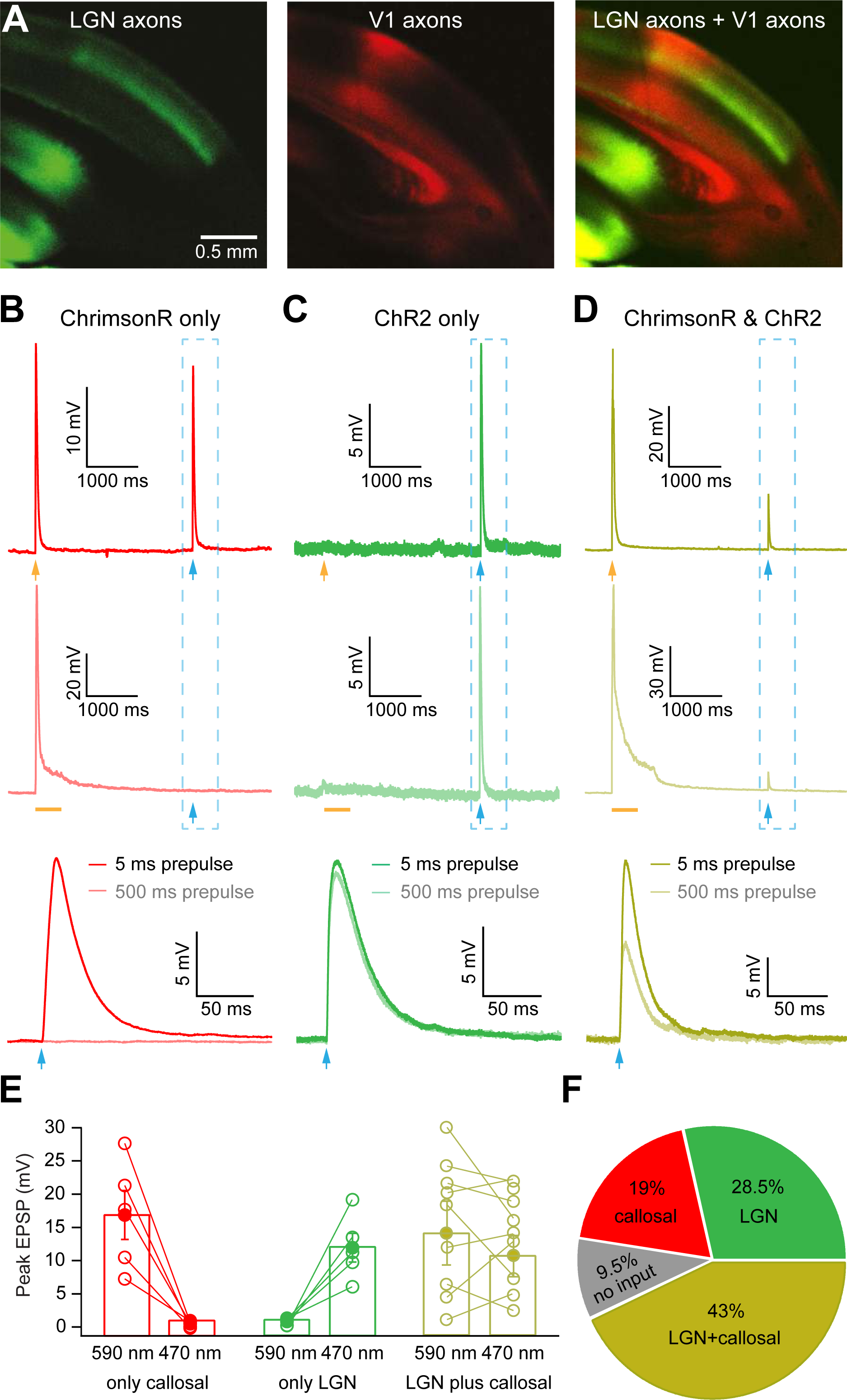
Convergence of callosal and ipsilateral lateral geniculate nucleus (LGN) input onto neurons in binocular V1. **A)** Coronal slices containing ChR2-eYFP axons from the ipsilateral LGN (left), ChrismonR-tdTomato callosal axons from the opposite V1 (middle), and an overlay showing convergence of LGN and callosal axons in binocular V1 (right). **B-D)** Examples of L2/3 pyramidal neurons in binocular V1 receiving only ChrimsonR callosal input (**B**), only ipsilateral ChR2 LGN input (**C**) and both ChrimsonR callosal and ChR2 LGN input (**D**). Top: Response of a L2/3 pyramidal neuron in binocular V1 to a brief (5 ms) 590 nm light pulse (orange arrow) followed by a brief (2 ms) 470 nm light pulse (blue arrow) in TTX plus 4-AP. Middle: Response of the same L2/3 pyramidal neuron to a prolonged (500 ms) 590 nm light pulse (orange bar) followed by a brief (2 ms) 470 nm light pulse (blue arrow). Bottom: Comparison of 470 nm light responses highlighted in the dashed grey boxes for L2/3 pyramidal neurons in binocular V1 receiving only ChrimsonR callosal input (**B**), only ipsilateral ChR2 LGN input (**C**) and both ChrimsonR callosal and ChR2 LGN input (**D**). **E)** Summary of synaptic response amplitude in L2/3 pyramidal neurons in binocular V1 receiving only ChrimsonR callosal input (red), only ipsilateral ChR2 LGN input (green) and both ChrimsonR callosal and ChR2 LGN input (blue) during brief (5 ms) 590 nm light pulses or a brief (2 ms) 470 nm light pulse which was preceded by a 500 ms 580 nm light pulse. **F)** Pie-chart showing the proportion of L2/3 pyramidal neurons in binocular V1 receiving direct, monosynaptic input from only contralateral V1 (red), only the LGN (green) or from both (blue). Cells not receiving LGN or contralateral V1 input are also indicated (grey).

While orange (590 nm) light will only activate ChrimsonR, due to overlap in the excitation spectra blue (470 nm) light will activate both ChR2 and ChrimsonR. Hence, responses evoked by blue light could be due to activation of ChR2-expressing axons from the ipsilateral LGN or ChrismonR-expressing callosal axons from contralateral V1. To get around this problem a prolonged pulse (500 ms) of orange light (590 nm) was used to reversibly inactivate ChrimsonR-expressing axons. To verify that this approach works, we recorded synaptic responses from L2/3 pyramidal neurons in binocular V1 in the presence of TTX plus 4-AP in animals injected with only ChrimsonR in contralateral V1. Pulses of orange (590 nm) light of varying duration (5 to 500 ms) were followed by a brief (2 ms) blue (470 nm) light pulse (Supplementary Figure 8A). As expected both orange and blue light pulses evoked synaptic responses in L2/3 pyramidal neurons receiving callosal input. Increasing the duration of the orange light pulse, however, reduced the amplitude of the synaptic response evoked by the blue light pulse, which was abolished when the preceding orange light pulse was 500 ms in duration (Supplementary Figure 8B,C). These data indicate that prolonged (500 ms) activation of ChrimsonR-expressing axons inactivates them temporarily (for ∼90 seconds; data not shown), such that they cannot be activated by a subsequent brief blue-light pulse. Finally, we verified in animals only injected with ChR2 in contralateral V1 that prolonged, 500 ms orange light pulses did not influence the amplitude of synaptic responses recorded in the opposite binocular V1 during blue light stimulation (Supplementary Figure 8D-F). In summary, these control experiments indicate that prolonged (500 ms) orange light pulses can be used to isolate ChrimsonR and ChR2-evoked synaptic responses despite the overlap in the excitation spectra of these two opsins.

Next, we made whole-cell current-clamp recordings from L2/3 pyramidal neurons in binocular V1 ipsilateral to the LGN ChR2 injection site in animals injected with both ChR2 in LGN and ChrismonR in contralateral V1. These experiments were performed in the presence of TTX plus 4-AP to isolate monosynaptic inputs. While callosal axons from the contralateral V1 will only be activated by orange (590 nm) light, to isolate ipsilateral LGN input blue light pulses were preceded by a long (500 ms) orange light pulse to temporarily inactivate ChrimsonR axons, as described above. L2/3 pyramidal neurons in binocular V1 could be classified into four distinct populations. One population received only callosal input, identified by responses to both orange and blue light, with the response to blue light abolished by a preceding (500 ms) orange light pulse (Fig. 8B). A second population received only LGN input, identified by a response to blue, but not orange, light (Fig. 8C). A third population received both callosal input from the contralateral V1 and ipsilateral LGN input, identified by responses to both orange and blue light, where the response to blue light was decreased, but not abolished by preceding (500 ms) orange light stimulation (Fig. 8D). Finally, a fourth population did not generate responses to either blue or orange light (not shown).

The synaptic response amplitudes for the above populations with and without a preceding orange light pulse are summarized in Fig. 8E, with the proportion of neurons in each population summarized in Fig. 8F. On average, the largest group, representing 43% of the population (9 out of 21 neurons), received monosynaptic input from both LGN and V1. The second largest group, representing 29% of the population (6 out of 21 neurons), received monosynaptic input only from the LGN. A third group, representing 19% of the population (4 out of 21 neurons), received monosynaptic input only from contralateral V1. A small percentage of neurons (9%; 2 out of 21) did not receive input from the LGN or contralateral V1. As neurons that received monosynaptic input from the contralateral V1 are binocular, whereas neurons that did not are monocular, these data indicate that the majority of binocular neurons receive direct, monosynaptic input from both the opposite V1 and the ipsilateral LGN.

Finally, we investigated the subcellular organization and spatial overlap of LGN and contralateral V1 inputs to binocular L2/3 pyramidal neurons. To do this, we recorded light-evoked synaptic responses in the presence of TTX plus 4-AP using LED stimulation with a 100 μm diameter spot to restrict illumination to specific parts of the dendritic tree. This 100 μm LED spot was incrementally moved in 50 μm steps along the basal/apical axis (Supplementary Fig 9A,B). LGN (ChR2) and contralateral V1 (ChrimsonR) synaptic responses at each location were isolated using the dual-color protocol outlined above (Supplementary Fig 8). These experiments indicated that the subcellular location of LGN and contralateral V1 callosal input onto pyramidal neurons is different, with significant overlap in basal dendrites, but with a preference for a more distal apical location of contralateral V1 callosal input (Supplementary Fig 9C-D).

## Discussion

The overall goal of this study was to identify the circuit pathways involved in binocular signal processing and to compare the cellular properties of binocular and monocular neurons. We show that by combining optogenetics and electrophysiology (*in vitro* and *in vivo*), we can identify binocular and monocular neurons by the presence or absence, respectively, of long-range callosal input from the opposite visual cortex. *In vivo* we found that ipsilateral eye responses in L2/3 pyramidal neurons in binocular V1 were significantly delayed compared to contralateral eye responses, suggesting that ipsilateral and contralateral eye inputs to V1 are mediated by different pathways. Since the majority of retinal axons cross at the optic chiasm in rodents (Drager and Olsen, 1980; Lund, 1965), we speculated that ipsilateral eye input arises primarily from the contralateral V1. Consistent with this idea, two-photon calcium imaging of callosal axons confirmed that callosal axons convey visual input from the ipsilateral eye to binocular V1. Furthermore, we show that optogenetic inhibition of the opposite V1 significantly reduces the ipsilateral, but not contralateral, synaptic eye response in binocular L2/3 pyramidal neurons. Finally, we show that only functionally identified binocular L2/3 pyramidal neurons received callosal input, whereas monocular neurons did not.

Using the presence of callosal input to identify binocular neurons *in vitro*, we show that binocular V1 contain two populations of L2/3 pyramidal neurons: one that receives callosal input and one that does not. At the cellular level, callosal receiving (binocular) L2/3 pyramidal neurons in binocular V1 had reduced excitability compared to monocular neurons, with the extent of this difference dependent on the magnitude of callosal input. Furthermore, we show that the reduced excitability of binocular neurons is due to a higher expression of Kv1 potassium channels. At the circuit level we show that only monocular, but not binocular, L2/3 pyramidal neurons in binocular V1 send a callosal projection to the opposite V1. Finally, we identify four populations of L2/3 pyramidal neurons in binocular V1 based on their inputs, finding that the largest population consists of binocular neurons that received monosynaptic input from both the ipsilateral LGN and contralateral V1.

Previous studies in rodents have provided contradictory evidence concerning the role of callosal projections to binocularity V1, with some concluding that the callosal pathway contributes ipsilateral eye information (Cerri et al., 2010; Diao et al., 1983; Drager, 1975; Restani et al., 2009; Zhao et al., 2013), whereas others concluding that it does not (Coleman et al., 2009; Hagihara et al., 2021; Liang et al., 2015; Olavarria, 2001; Ramachandra et al., 2020). Consistent with our findings, previous studies using extracellular recording and pharmacological inactivation of contralateral V1 with muscimol (Restani et al., 2009) or TTX (Zhao et al., 2013), support the idea that callosal projections carry ipsilateral eye information. In addition, recent *in vivo* 2-photon imaging of individual spines on L2/3 pyramidal neurons in binocular V1 indicates that spines receiving callosal inputs have a strong bias for the ipsilateral eye (Lee et al., 2019). Our data, using whole-cell recording to measure synaptic responses, reinforces this idea and provides strong evidence at a subthreshold level that interhemispheric projections convey ipsilateral eye input directly to binocular L2/3 pyramidal neurons in binocular V1.

The fact that ipsilateral eye responses were not abolished in some neurons by inhibition of the opposite V1 may indicate only partial optogenetic inhibition of the opposite V1, or could suggest that a component of the ipsilateral eye response is mediated via the small percentage of uncrossed RGC axons projecting to the LGN on the same side (Coleman et al., 2009; Grieve, 2005; Hagihara et al., 2021; Howarth et al., 2014; Laing et al., 2015; Lewis and Olavarria, 1995; Olavarria and Van Sluyters, 1983; Olavarria, 2001; Ramachandra et al., 2020). While we cannot distinguish between these two possibilities, our finding that the amplitude of ipsilateral eye responses is correlated with the magnitude of callosal input, together with our observation of an almost one-to-one correlation between ipsilateral eye responses and optogenetic responses to activation of callosal input, suggest that ipsilateral eye input is primarily conveyed to binocular neurons via the callosal pathway. The alternative explanation of these findings would be that ipsilateral eye input to binocular neurons from the callosal projection and ipsilateral LGN may be correlated, with neurons that respond strongly to the ipsilateral eye receiving strong input from both pathways.

Our findings are inconsistent with two recent studies (Hagihara et al., 2021; Ramachandra et al., 2020). Hagihara *et al*. (2021) found in mouse visual cortex that callosal projecting (monocular) neurons have a bias for the ipsilateral eye, whereas Ramachandra *et al*. (2020) found that roughly equal proportions of binocular and monocular neurons contribute callosal projections to the opposite V1 in rats. There are a number of possible explanations for the different conclusions made by these studies and ours. Firstly, both of these studies classify callosal projection neurons based on visual inspection of retrograde labeling *in vivo*, which could lead to errors in classification of projecting and non-projecting neurons. We overcome this potential bias by retrograde expression of ChR2, allowing us to identify callosal projecting neurons post-hoc based on their electrical response to excitation of ChR2. Secondly, *in vivo* 2-photon calcium imaging as used by Hagihara *et al*. (2021) and Ramachandra *et al*. (2020) may lead to errors in the functional classification of binocular and monocular neurons for a number of reasons. Firstly, in contrast to whole-cell patch-clamp, 2-photon calcium imaging gives an indirect readout of electrical activity and only detects suprathreshold spiking activity. As a result, binocular cells with subthreshold ipsilateral eye responses would be classified as monocular. Secondly, ocular stimulation of the ipsilateral (and contralateral) eye can cause delayed bursts of action potentials, similar to up-states. Given the low time resolution of 2-photon calcium imaging it would be difficult to distinguish these events from visual responses due to direct eye stimulation. As a result, some monocular cells may have been incorrectly classified as binocular by Hagihara *et al*. (2021) and Ramachandra *et al*. (2020). In contrast, our classification is based on measurement of subthreshold synaptic responses at high time resolution using whole-cell recording, allowing us to unequivocally identify neurons as binocular or monocular based on their synaptic input.

At the cellular level, we found that layer 2/3 binocular neurons were significantly less excitable that monocular neurons despite no obvious differences in passive or morphological properties. Furthermore, this difference in excitability was correlated with the amplitude of callosal input. Consistent this *in vitro* data, the excitability of functionally identified binocular neurons *in vivo* was also correlated with the amplitude of the ipsilateral eye response. The reduced intrinsic excitability of binocular compared to monocular neurons was due to a higher expression of D-type Kv1 potassium channels. The finding that Kv1/D-type potassium channels play a critical role in regulating neuronal excitability is consistent with previous studies (Bekkers and Delaney, 2001; Metz et al., 2007; Ordemann et al., 2019). Given that previous studies have shown that ion channel expression is activity dependent and regulates homeostatic mechanisms (Turrigiano and Nelson, 2004), our findings suggest that the reduced intrinsic excitability of binocular neurons may result from the increased excitatory drive binocular neurons receive via callosal input. Consistent with this idea, we found that the excitability of binocular neurons was negatively correlated with the strength of both callosal and ipsilateral eye input. These data are consistent with recent studies which suggest that L2/3 pyramidal neurons in the superficial layers of binocular V1 are organized as a continuum and show a gradient in terms of their molecular (Cheng et al., 2022; Kim et al., 2020; Tan et al., 2020) and electrophysiological properties (Weiler et al., 2023).

At the circuit level, we identified both optically and electrically that monocular neurons send callosal projections, but binocular neurons do not. This observation is consistent with the idea that monocular and binocular neurons represent distinct populations within the binocular band of V1. Furthermore, it suggests that callosal projections carry solely ipsilateral eye information. Consistent with this idea, optogenetic inhibition of the opposite V1 reduced ipsilateral, but not the contralateral, eye responses in binocular L2/3 neurons. Together, our observations suggest that only monocular neurons send callosal projections, whereas only binocular neurons receive callosal input. We propose that this reciprocal binocular circuit “logic” (shown schematically in Supplementary Fig. 10) makes intuitive sense and is likely to be conserved across species. Further, we hypothesize it is likely to play a key role in the generation of a cohesive percept despite binocular information being processed by both hemispheres independently. Future work will be required to determine the molecular mechanisms underlying this circuit motif as well as the role of other circuit components, such as different inhibitory neuronal populations.

To conclude, using whole-cell recording and optogenetics *in vivo* and *in vitro* we identify a critical role of the callosal input from the opposite V1 in binocular visual processing in rodents. At the cellular level we find that binocular L2/3 pyramidal neurons are less excitable than monocular neurons due to higher expression of Kv1 potassium channels. At the circuit level we find that only binocular L2/3 pyramidal neurons receive callosal input from the opposite V1, whereas only monocular L2/3 pyramidal neurons send callosal projections to the opposite V1. Finally, using dual-colour optogenetics we show that binocular neurons receive input from both the opposite V1 and the LGN, whereas monocular neurons receive input only from the LGN. These experiments provide crucial insights into the cellular and circuit mechanisms underlying the processing and integration of binocular visual information.

## Acknowledgments

We thank Gord Fishell for comments on the manuscript. This work was funded via support from the Australian National University, Monash University, the National Health and Medical Research Council of Australia and the Australian Research Council Centre of Excellence for Integrative Brain Function.

---

**Supplementary Figure 1.**
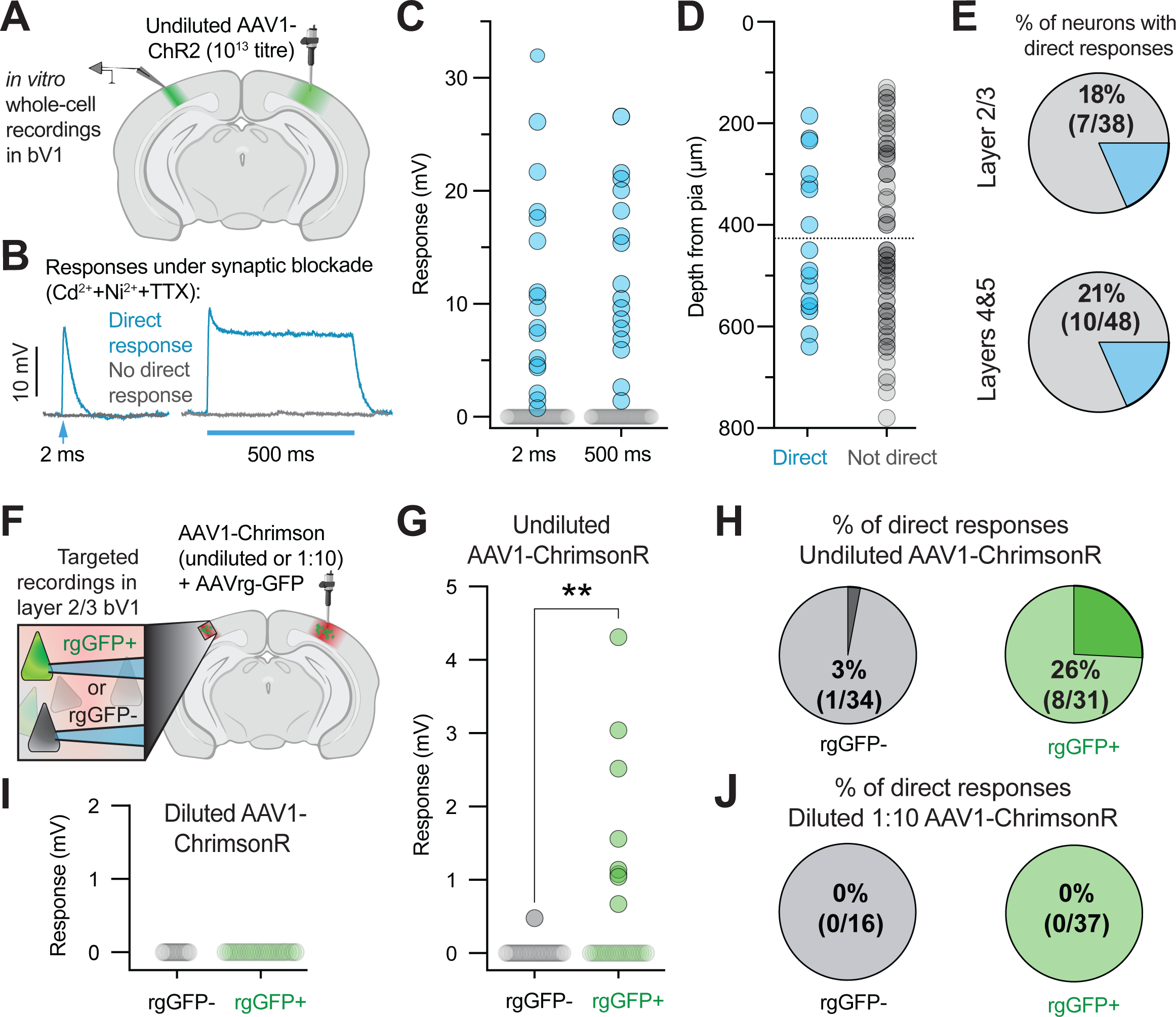
Dilution of Addgene AAV1 abolishes retrograde transport. **A)** Schematic of experiments **(B-E)**. Undiluted AAV1-ChR2 (Addgene; 10^13^ vg/mL titre) was injected into right binocular V1 (bV1) of adult mice (n=4 animals). 3-4 weeks later *in vitro* whole-cell recordings were made from left bV1. **B)** Responses evoked by brief (left; 2 ms) and long (right; 500 ms) light pulses (470 nm, 4 mW) under synaptic blockade of transmitter release with cadmium chloride (Cd^2+^, 200 µM) plus nickel chloride (Ni^2+^,1 mM) and tetrodotoxin (TTX, 500 nM) in neurons with (blue) and without a direct ChR2 response (grey). **C)** Average amplitude of light-evoked responses (2 ms) recorded under the same conditions as **(B)** in neurons with (blue) and without (grey) a direct ChR2 response. **D)** Depth from pia of neurons shown in **(C)** in neurons with (blue) and without (grey) a direct ChR2 response. **E)** Proportion of neurons with direct responses in layer 2/3 (top) and layers 4&5 (bottom). **F)** Schematic of experiments **(G-J)**. Undiluted AAV1-ChrimsonR (Addgene; 10^13^ vg/mL titre) or diluted (1:10 in saline) AAV1-ChrimsonR (Addgene; 10^12^ vg/mL titre) was co-injected with retrograde GFP (rgGFP) into right bV1. 3-4 weeks later targeted *in vitro* whole-cell recordings were made from rgGFP+ and nearby rgGFP-neuron in layer 2/3 of left bV1. In **(G,H)** undiluted AAV1-ChrimsonR was used (n=4 mice), whereas in **(I,J)** diluted AAV1-Chrimson was used (n=3 mice). **G,I)** Average ampltidue of light-evoked responses (2 ms) under synaptic blockade of glutamate and GABA receptors with DNQX (20 µM), APV (25 µM) and gabazine (10 µM) in rgGFP+ (green) and rgGFP-(grey) neurons. Responses in rgGFP+ neurons were significantly larger than in rgGFP-neurons in mice injected with undiluted AAV1-Chrimson R (Two-tailed t-test: P<0.01), but absent in both rgGFP+ and rgGFP-neurons in mice inject-ed with diluted AAV1-Chrimson. **H,J)** Proportion of rgGFP-(left) and rgGFP+ (right) neurons with direct responses in mice injected with undiluted (**H**) and diluted (**J**) AAV1-ChrimsonR.

**Supplementary Figure 2:**
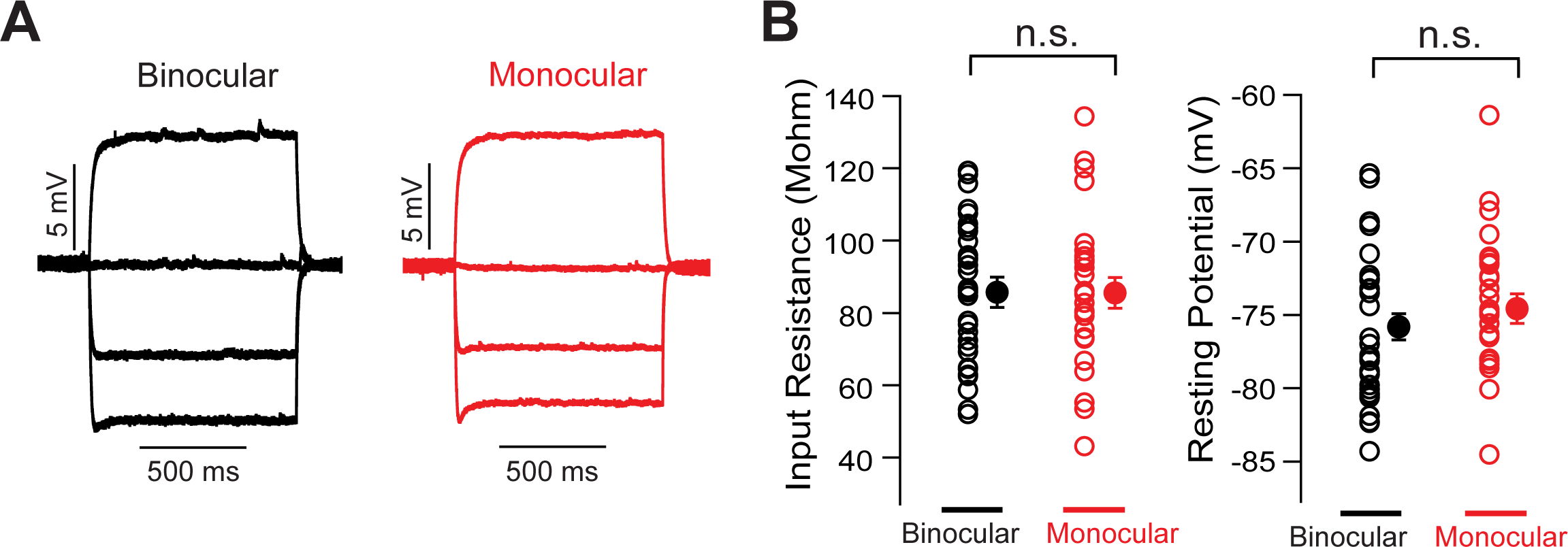
Passive properties of binocular and monocular neurons are similar. **A)** Subthreshold voltage responses in layer 2/3 pyramidal neurons during somatic current injection (−200, −100, 0 and +100 pA) in binocular (black) and monocular (red) neurons. **B)** Pooled data from binocular (black) and monocular (red) neurons showing individual and mean input resistance and resting membrane potential in these neuronal populations (n=59). Error bars represent s.e.m and n.s indicates no significant difference (P>0.05). Cells were identified as binocular or monocular based on the presence or absence of callosal input.

**Supplementary Figure 3:**
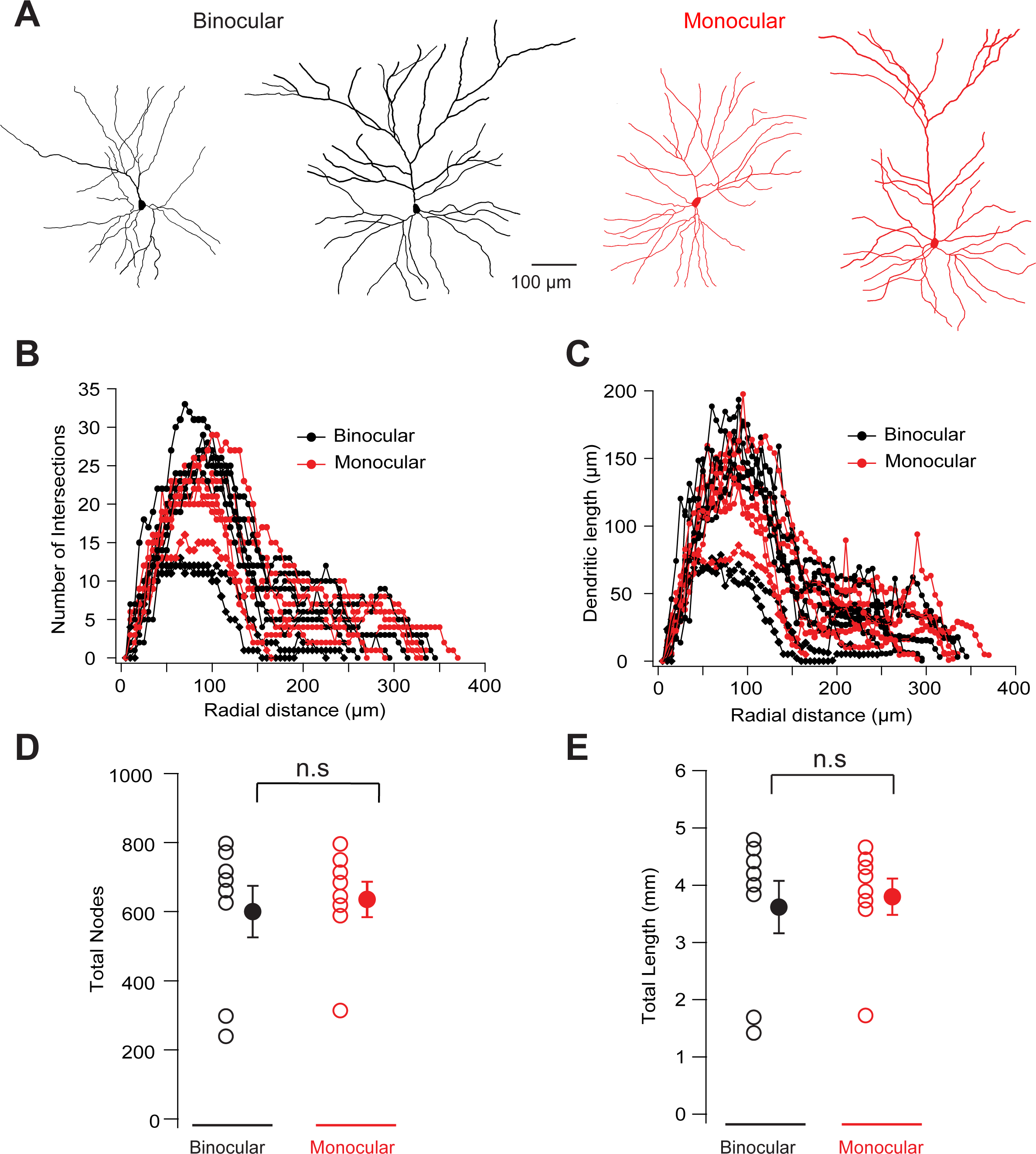
Morphological properties of binocular and monocular neurons are not different. **A)** Morphological reconstructions of biocytin filled binocular and monocular L2/3 pyramidal neurons from superficial (∼200 μm from pia) and deeper (∼350 μm from pia) binocualr V1. **B,C)** Quantification of dendritic branching and geometry using Scholl analysis showing the number of intersections **(B)** and dendritic length **(C)** as a function of radial distance from soma. **D,E)** Pooled data showing the total number of nodes **(D)** and total dendritic length **(E)** in different binocular and monocular neurons. Error bars represent s.e.m and n.s indicates no significant difference (P>0.05). Cells were identified as binocular or monocular based on the presence or absence of callosal input.

**Supplementary Figure 4.**
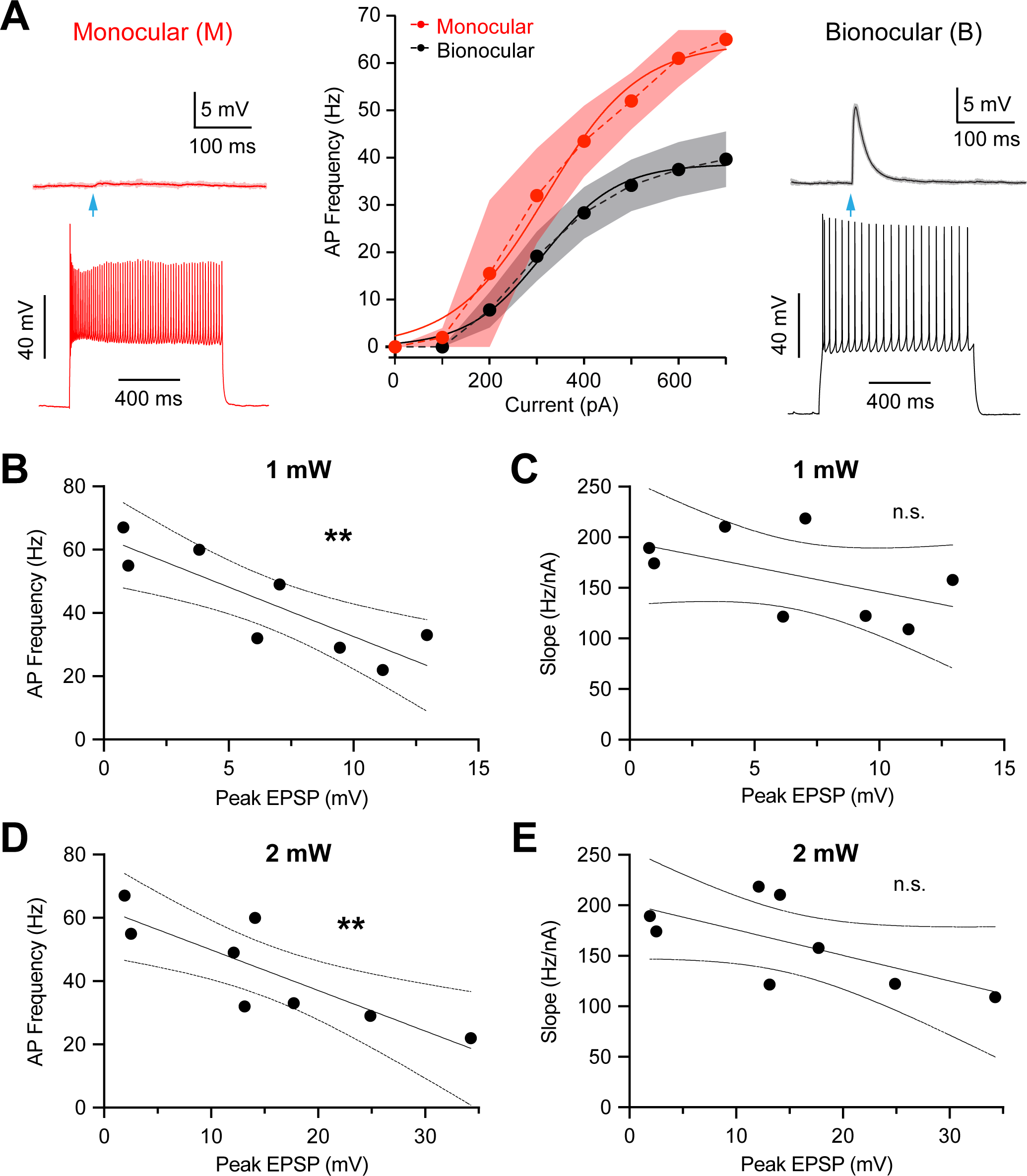
Callosal input and excitability in mice injected with 1:10 diluted AAV1-ChR2. Data from animals injected with 1:10 diluted AAV1-ChR2 in the opposite V1 (n=5 mice, n=8 neurons). **A)** Left & Right: Synaptic responses (top; 3 trials; individual trials shaded, average bold) evoked by brief (2 ms, 470 nm, 1 mW) light pulses (blue arrows) and action potential firing evoked by somatic current injection (bottom; 600 pA, 1 s) in a monocular (left) and binocular (right) layer 2/3 neuron. Middle: Pooled AP firing frequency in response to depolarising current injections (1 s, 0-700 pA) in monocular (n=2) and binocular (n=6) layer 2/3 neurons. Solid lines represent sigmoid fit ± SEM (shad-ing). **B-E)** Plot of callosal EPSP amplitude during optogenetic activation at 1 mW (**B,C**) or 2 mW (**D,E**) versus AP firing frequency for a 600 pA somatic current pulse (**B,D**) and sigmoid slope (**C,E**) for differ-ent neurons (n=8). Callosal EPSP amplitude and AP firing frequency (**B**,**D**) were significantly correlat-ed at both LED powers (1 mW: R^2^ = 0.73, P<0.01; 2 mW: R^2^ = 0.71, P<0.01), whereas callosal EPSP amplitude and f/I slope (**C**,**E**) were not (P>0.05). Pearson correlation (solid line) ± 95% confidence interval (dotted lines).

**Supplementary Figure 5.**
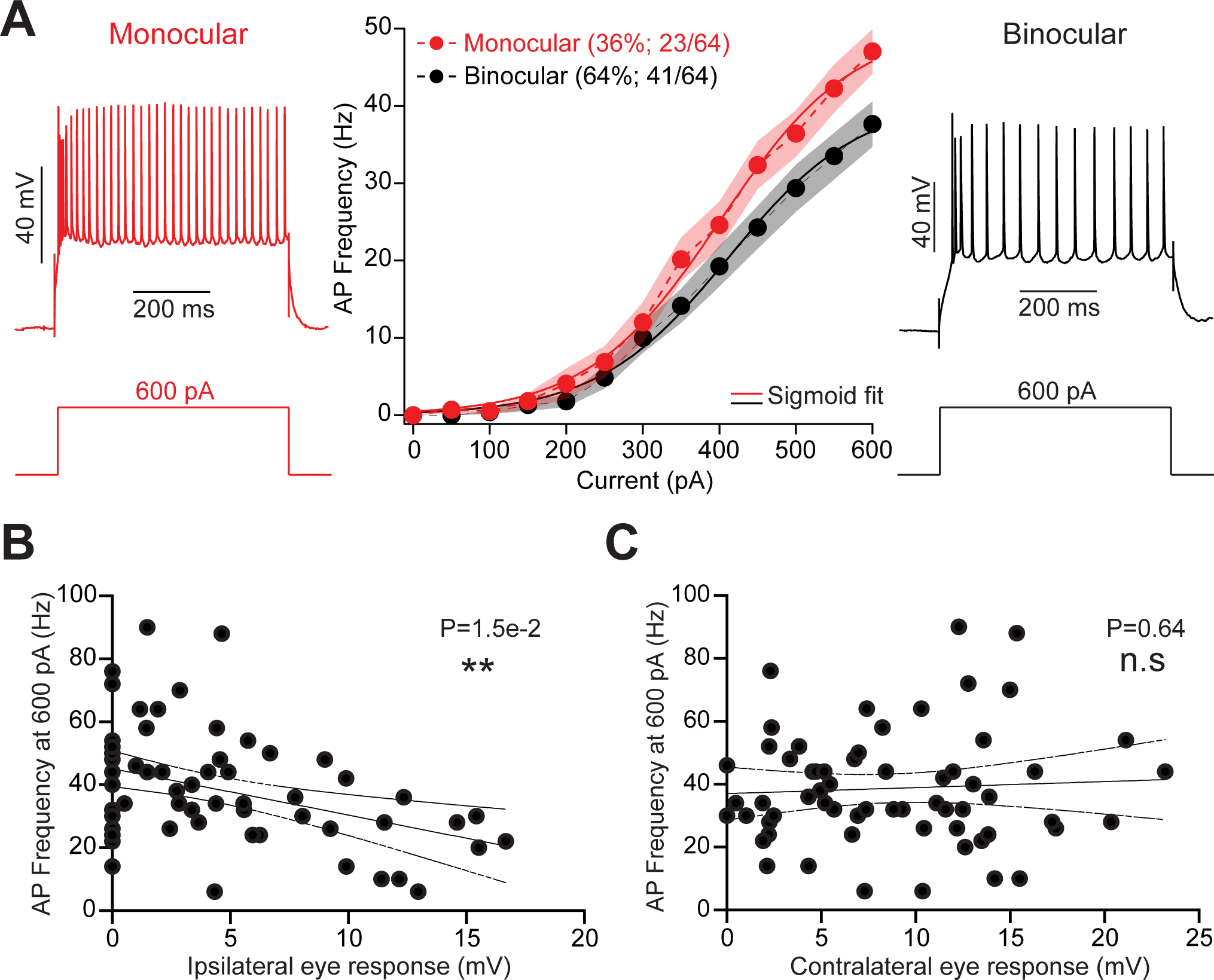
Excitability of monocular versus binocular neurons *in vivo*. **A)** Action potential (AP) response (top) to 600 pA current pulse (bottom) in functionally identified monocular (left) and binocular (right) neurons in binocular visual cortex *in vivo*. Middle, Average action potential frequency versus current (f/I) relationship for monocular (red) and binocular (black) neurons. Solid line sigmoid fit; Shading SEM. **(B,C)** Relationship between AP firing frequency evoked by 600 pA curent injections and the amplitude of the ipsilateral **(B)** and contra-lateral **(C)** eye response. Error bars represent s.e.m, n.s indicates no significant difference (P>0.05) and asterisks indicate P<0.05.

**Supplementary Figure 6:**
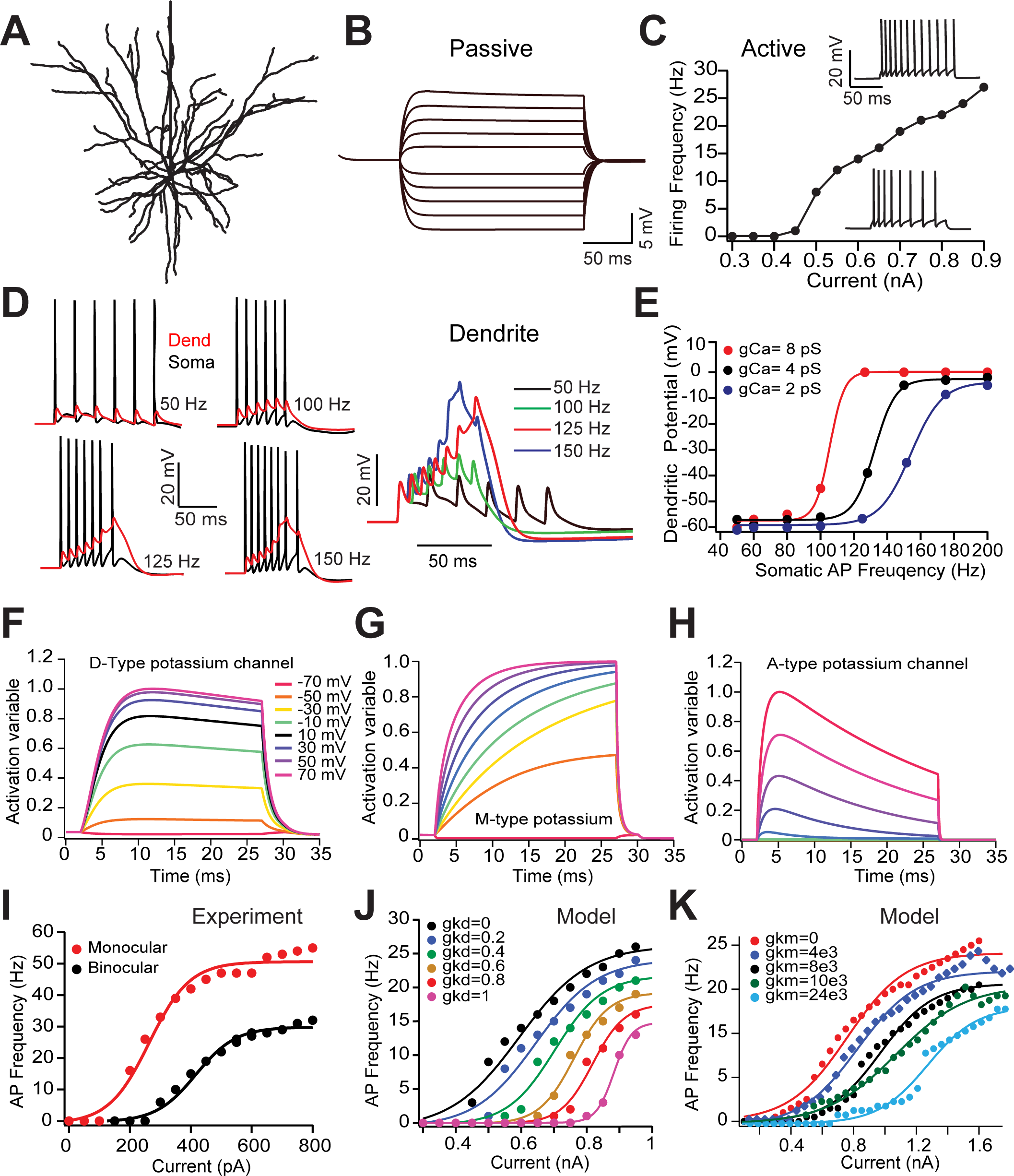
Impact of different potassium channels on f/I curves in a morphologically realistic layer 2/3 neuron model. **A)** Morphology of a reconstructed layer 2/3 pyramidal neuron obtained from NeuroMorpho.org ID Martin, NMO_00904 (Smith et al., 2013). **B)** Somatic response of the model to somatic current injection (−300 to +200 pA in 50 pA steps). **C)** Action potential firing frequency of the layer 2/3 neuron model during somatic current injection from (+300 to +900 pA). Inserts show action potential firing during 500 pA (bottom) and 800 pA (top) current steps. **D)** Tuning dendritic voltage-activated Ca2+ channel density to match dendritic elec-trogenesis evoked by trains of backpropagating action potentials at different frequencies to match experi-mental data (Larkum et al., 2006). **E)** Critical frequency plots for different dendritic voltage-activated Ca2+ channel densities. 2 pS (blue), 4 pS (black) and 8 pS (red). **F-H)** Activation variable kinetics for voltage-acti-vated D-type (Kv1.1/2; **F**), M-type (Kv7.2/3; **G**) and A-type (Kv4.2; **H**) K+ channels obtained from a single compartment model. **I)** Experimental data showing experimental differences in the f/I curve in binocular (black) and monocular (red) layer 2/3 pyramidal neurons. **J)** Graded impact of different axonal Kv1.1 chan-nel conductance densities (gKd) on the f/I curve in the layer 2/3 neuron model. **K)** Graded impact of different axonal Kv7.2 channel conductance densities (gKm) on the f/I curve in the layer 2/3 neuron model.

**Supplementary Figure 7.**
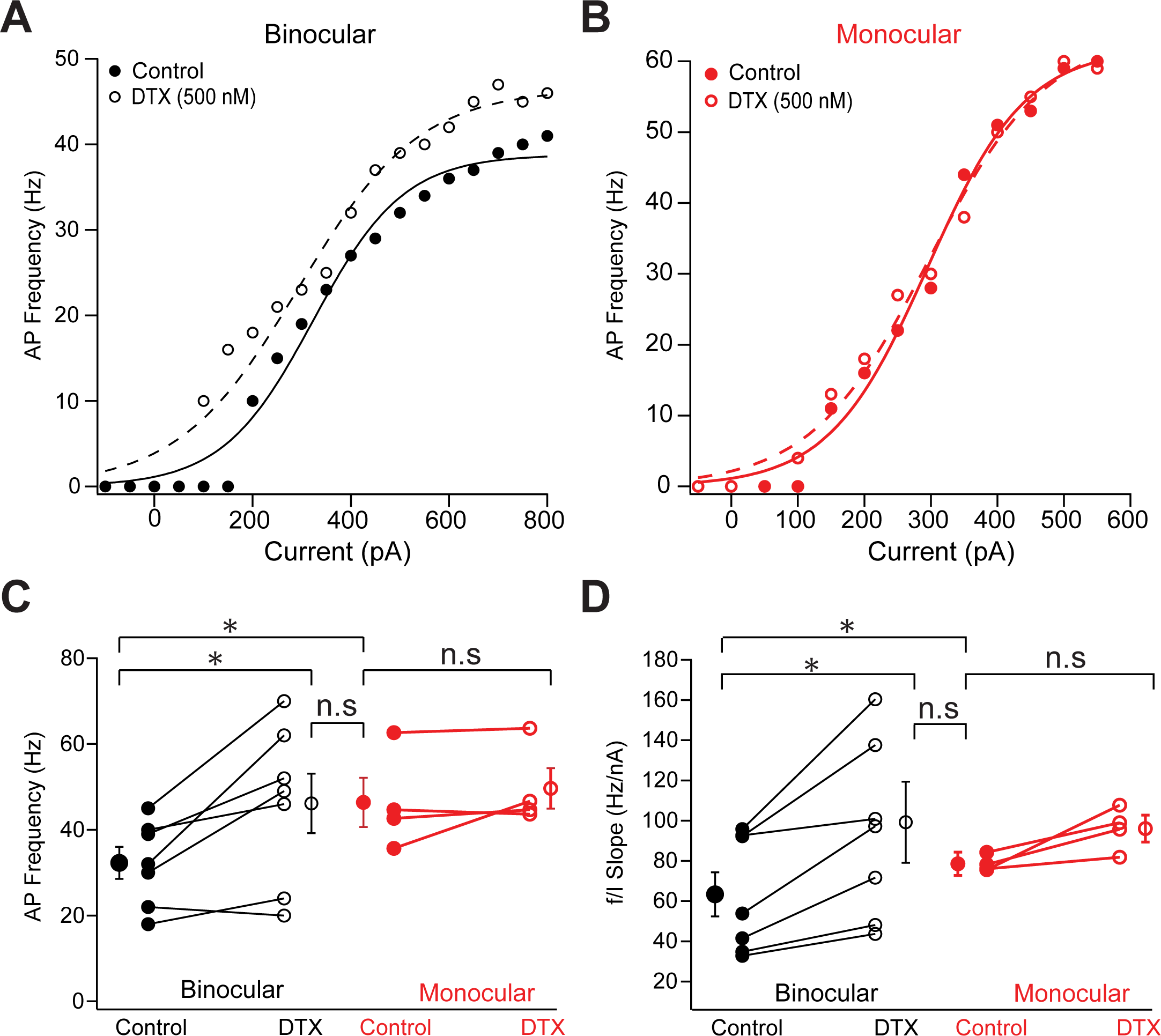
Differences in the excitability of binocular and monocular neu-rons are abolished by dendrotoxin (DTX) **A,B)** Action potential (AP) firing frequency versus current injection (f/I) curves in control (closed circles) and after application of DTX (open circles) in putative binocular (**A**, black) and monocular neurons (**B**, red). **C,D)** Maximum AP firing rate to a 600 pA current pulse **(D)** and f/I slope obtained from sigmoid fits to data like that shown in **(A,B)** in control and after application of DTX (500 nM) in binocular (black) and monocular (red) neurons. Error bars represent s.e.m and n.s indicates no significant difference (P>0.05). Asterisks indicate significant differences using a Wilcoxon Mann Whitney signed ranked test (P < 0:05 (*)).

**Supplementary Figure 8.**
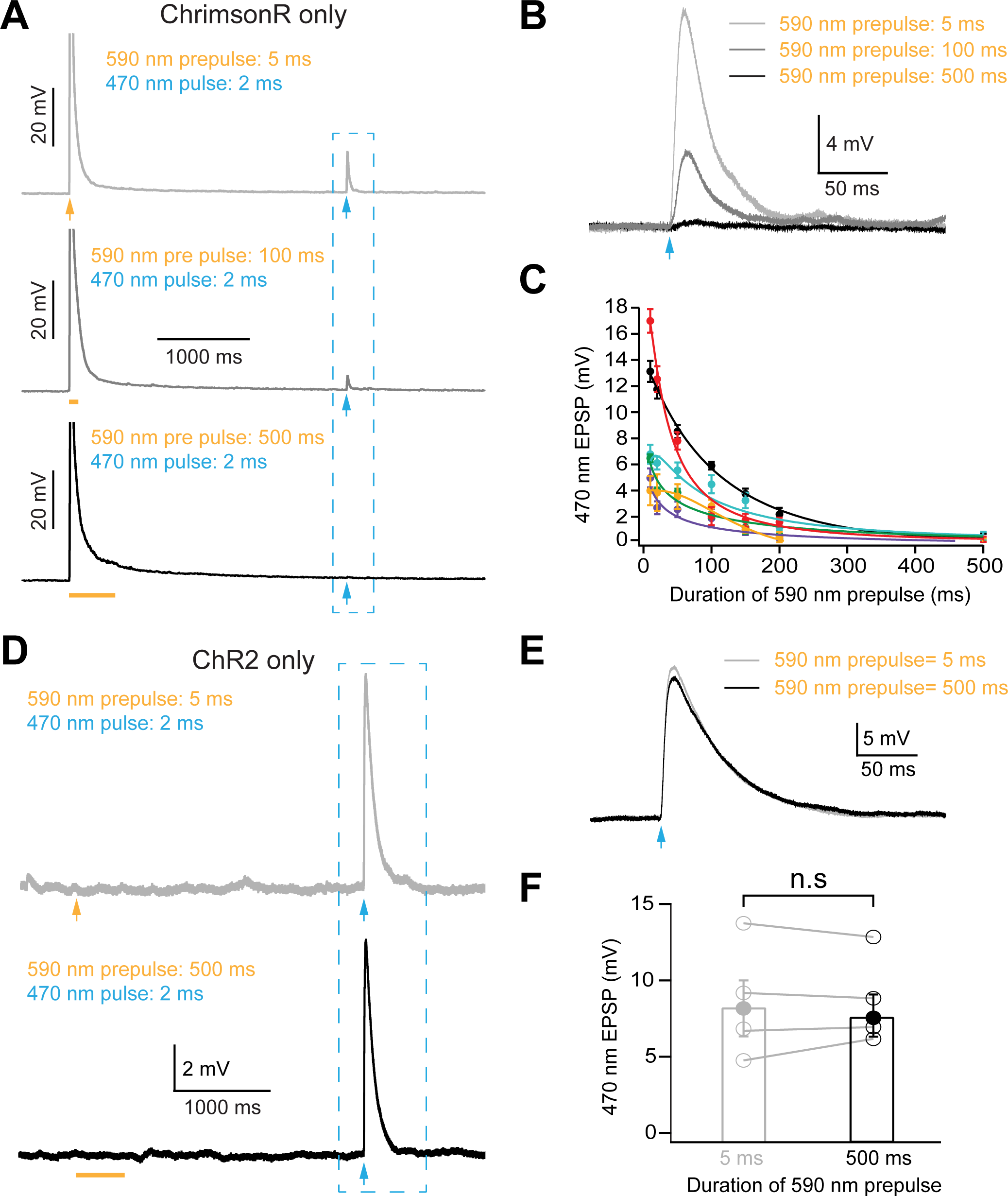
Isolation of dual wavelength optogenetic responses. **A)** Synaptic responses evoked by brief (2 ms) 470 nm light pulses (blue arrows) preceeded by 5 ms (top; orange arrow), 100 ms (middle; orange bar) or 500 ms (bottom; orange bar) 590 nm light pulses in a layer 2/3 binocular neuron expressing only ChrimsonR in callosal axons. **B)** Expan-sion of the responses shown in the blue dashed box in panel (**A**). **C)** Impact of the duration of preceding 590 nm light pulse on the amplitude of responses to a brief (2 ms) 470 nm light pulse in different neurons (different colours) in animals expressing only ChrimsonR in callosal axons. **D)** Synaptic responses evoked by a brief (2 ms) 470 nm light pulse (blue arrows) preceeded by a 5 ms (top; orange arrow) or 500 ms (bottom; orange bar) 590 nm light pulse in a layer 2/3 binocular neuron expressing only ChR2 in callosal axons. **E)** Expansion of the responses shown in the blue dashed box in panel (**D**). **F)** Summary data showing the impact of a preceding brief (5 ms) or prolonged (500 ms) 590 nm light pulse on synaptic responses evoked by a brief (2 ms) 470 nm light pulse in animals expressing only ChR2 in callosal axons.

**Supplementary Figure 9:**
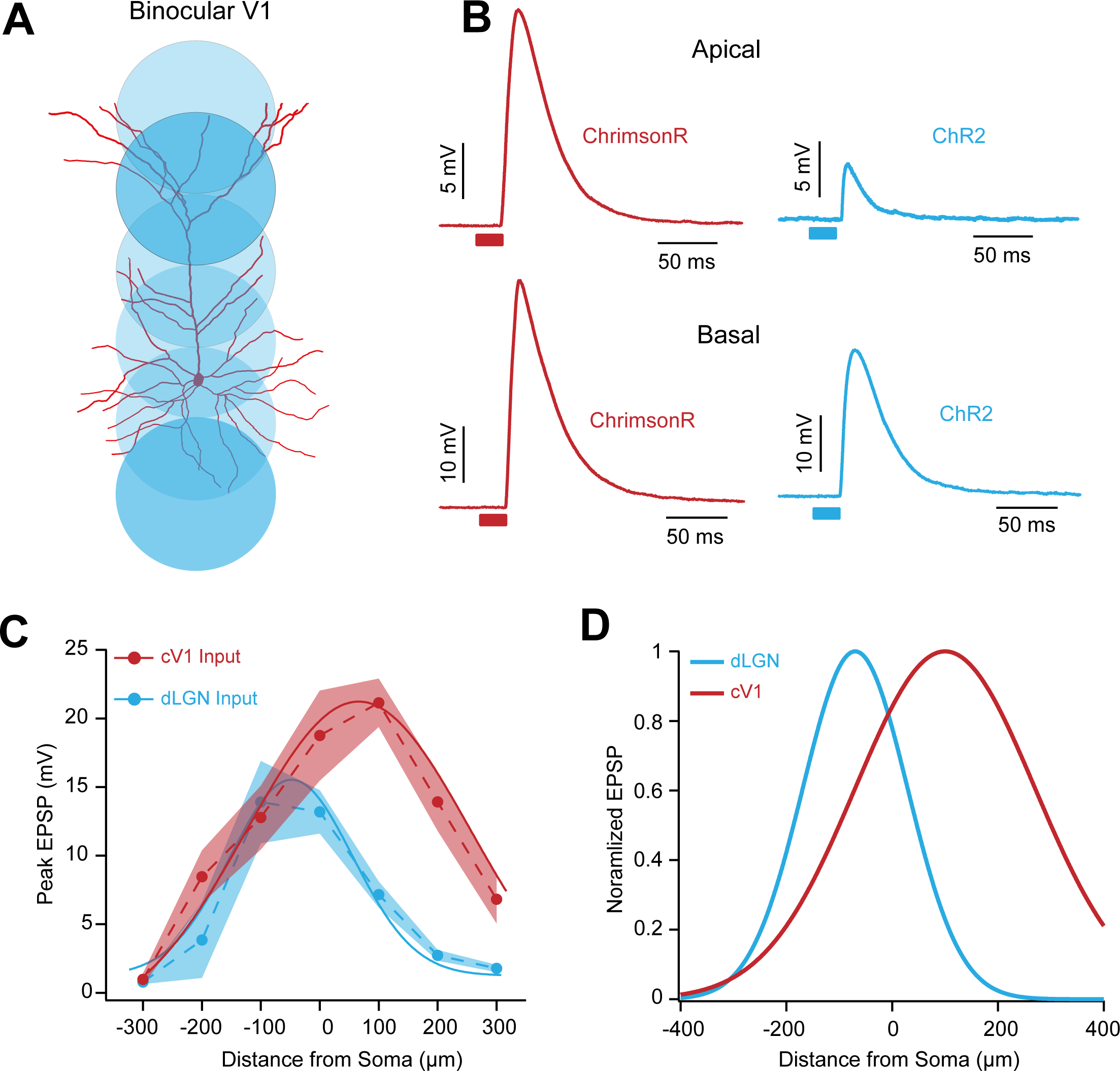
Subcellular mapping of LGN and contralateral V1 synaptic input to bin-ocular V1 neurons. **A)** Schematic of the experimental paradigm showing restricted LED 470 nm (blue) illumination spots (100 μm diameter spots placed at 50 μm intervals along the basal-apical axis) superimposed on a layer 2/3 pyramidal neuron. **B)** Representative isolated ChrimsonR (red) and ChR2 (blue) synaptic responses in a layer 2/3 pyramidal neuron during restricted 470 and 590 nm LED illumination of basal (bottom) and apical (top) dendrites of a binocular layer 2/3 pyramidal neuron following injection of ChR2 into ipsilater-al LGN and ChrimsonR into contralateral V1. ChR2 responses were preceded by 500 ms 590 nm light-pulses. **C)** Summary data showing averaged synaptic response amplitude (± SEM) of isolated ChrimsonR (red) and ChR2 (blue) responses, reflecting contralateral V1 and LGN input, generated by 100 μm LED spots positioned at different distances from the soma along the basal-apical axis (n=3; dotted lines). Data fitted with a gaussian. **D)** Normalized EPSP response versus distance of the LED spot from the soma quantifying functional overlap of LGN (blue) and contralateral V1 (red) inputs to binocular layer 2/3 binocular pyramidal neurons. Data fitted with a gaussian.

**Supplementary Figure 10:**
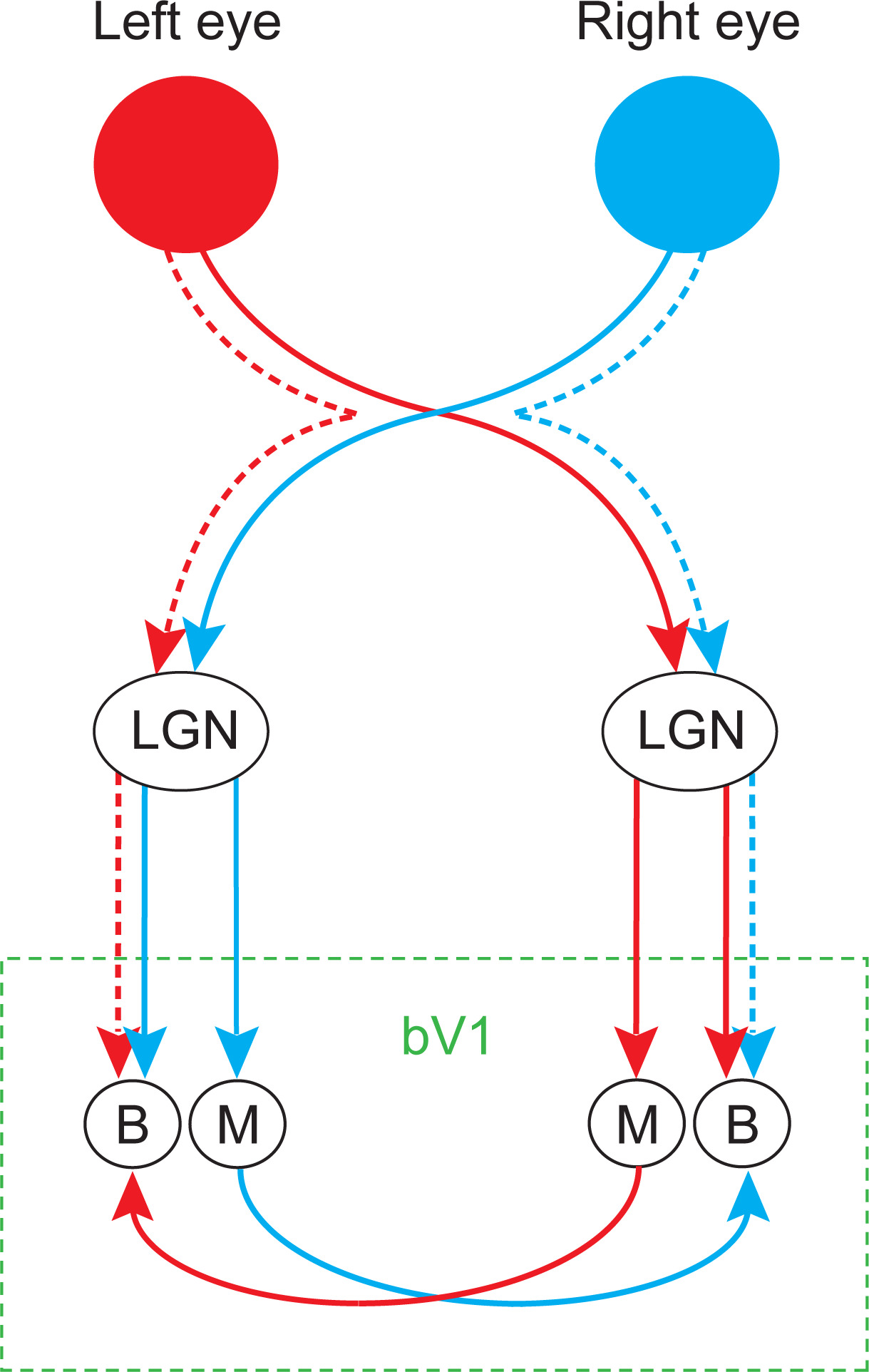
The binocular circuit. Schematic of the potential binocular circuit. Retinal ganglion cell axons from the left (red) and right (blue) eye project to the lateral genicular nucleus (LGN) in both hemispheres. The LGN then proj-ects to both monocular (M) and binocular (B) neurons in binocular primary visual cortex (bV1; green rectangle). Only binocular (B) neurons recieve callosal input from the opposite bV1, whereas only monocular (M) neurons send callosal projections to the opposite bV1. Solid lines represent the pathways determined in the current study. Dotten lines represent the uncrossed pathway.

## REFERENCES

Atallah, B.V., Bruns, W., Carandini, M., and Scanziani, M. (2012). Parvalbumin-expressing interneurons linearly transform cortical responses to visual stimuli. Neuron 73, 159–170.

Barlow, H.B., Blakemore, C., and Pettigrew, J.D. (1967). The neural mechanism of binocular depth discrimination. J Physiol 193, 327–342.

Battefeld, A., Tran, B.T., Gavrilis, J., Cooper, E.C., and Kole, M.H. (2014). Heteromeric Kv7.2/7.3 channels differentially regulate action potential initiation and conduction in neocortical myelinated axons. J Neurosci 34, 3719–3732.

Bauer, J., Weiler, S., Fernholz, M.H.P., Laubender, D., Scheuss, V., Hubener, M., Bonhoeffer, T., and Rose, T. (2021). Limited functional convergence of eye-specific inputs in the retinogeniculate pathway of the mouse. Neuron 109, 2457–2468 e2412.

Bekkers, J.M. (2000). Distribution and activation of voltage-gated potassium channels in cell-attached and outside-out patches from large layer 5 cortical pyramidal neurons of the rat. J Physiol 525 Pt 3, 611–620.

Bekkers, J.M., and Delaney, A.J. (2001). Modulation of excitability by alpha-dendrotoxin-sensitive potassium channels in neocortical pyramidal neurons. J Neurosci 21, 6553–6560.

Berlucchi, G., and Rizzolatti, G. (1968). Binocularly Driven Neurons in Visual Cortex of Split-Chiasm Cats. Science 159, 308-+.

Blakemore, C., Diao, Y., Pu, M., Wang, Y., and Xiao, Y. (1983). Possible Functions of the Interhemispheric Connections between Visual Cortical Areas in the Cat. Journal of Physiology-London 337, 331–349.

Brainard, D.H. (1997). The Psychophysics Toolbox. Spat Vis 10, 433–436.

Carnevale, N., and Hines, M. (2006). The NEURON Book. Cambridge University Press.

Cerri, C., Restani, L., and Caleo, M. (2010). Callosal contribution to ocular dominance in rat primary visual cortex. Eur J Neurosci 32, 1163–1169.

Chen, T.W., Wardill, T.J., Sun, Y., Pulver, S.R., Renninger, S.L., Baohan, A., Schreiter, E.R., Kerr, R.A., Orger, M.B., Jayaraman, V., et al. (2013). Ultrasensitive fluorescent proteins for imaging neuronal activity. Nature 499, 295–300.

Cheng, S., Butrus, S., Tan, L., Xu, R., Sagireddy, S., Trachtenberg, J.T., Shekhar, K., and Zipursky, S.L. (2022). Vision-dependent specification of cell types and function in the developing cortex. Cell 185, 311–327 e324.

Coleman, J.E., Law, K., and Bear, M.F. (2009). Anatomical origins of ocular dominance in mouse primary visual cortex. Neuroscience 161, 561–571.

Cusick, C.G., and Lund, R.D. (1981). The distribution of the callosal projection to the occipital visual cortex in rats and mice. Brain Res 214, 239–259.

Dehmel, S., and Lowel, S. (2014). Cortico-cortical interactions influence binocularity of the primary visual cortex of adult mice. PLoS One 9, e105745.

Diao, Y.C., Wang, Y.K., and Pu, M.L. (1983). Binocular responses of cortical cells and the callosal projection in the albino rat. Exp Brain Res 49, 410–418.

Drager, U.C. (1975). Receptive fields of single cells and topography in mouse visual cortex. J Comp Neurol 160, 269–290.

Drager, U.C., and Olsen, J.F. (1980). Origins of crossed and uncrossed retinal projections in pigmented and albino mice. J Comp Neurol 191, 383–412.

Elberger, A.J., and Smith, E.L., 3rd (1985). The critical period for corpus callosum section to affect cortical binocularity. Exp Brain Res 57, 213–223.

Gharaei, S., Honnuraiah, S., Arabzadeh, E., and Stuart, G.J. (2020). Superior colliculus modulates cortical coding of somatosensory information. Nat Commun 11, 1693.

Gordon, J.A., and Stryker, M.P. (1996). Experience-dependent plasticity of binocular responses in the primary visual cortex of the mouse. J Neurosci 16, 3274–3286.

Grieve, K.L. (2005). Binocular visual responses in cells of the rat dLGN. J Physiol 566, 119–124.

Hagihara, K., Ishikawa, A., Yoshimura, Y., Tagawa, Y., and Ohki, K. (2021). Long-range interhemispheric projection neurons show biased response properties and fine-scale local subnetworks in mouse visual cortex. Cerebal Cortex 31, 1307–1315.

Harvey, A.L. (2001). Twenty years of dendrotoxins. Toxicon 39, 15–26.

Hooks, B.M., Lin, J.Y., Guo, C., and Svoboda, K. (2015). Dual-channel circuit mapping reveals sensorimotor convergence in the primary motor cortex. J Neurosci 35, 4418–4426.

Howarth, M., Walmsley, L., and Brown, T.M. (2014). Binocular integration in the mouse lateral geniculate nuclei. Curr Biol 24, 1241–1247.

Hubel, D.H., and Wiesel, T.N. (1962). Receptive fields, binocular interaction and functional architecture in the cat’s visual cortex. J Physiol 160, 106–154.

Jacobson, S. (1970). Distribution of commissural axon terminals in the rat neocortex. Exp Neurol 28, 193–205.

Kennedy, H., Dehay, C., and Bullier, J. (1986). Organization of the Callosal Connections of Visual Areas V1 and V2 in the Macaque Monkey. Journal of Comparative Neurology 247, 398–415.

Kim, E.J., Zhang, Z., Huang, L., Ito-Cole, T., Jacobs, M.W., Juavinett, A.L., Senturk, G., Hu, M., Ku, M., Ecker, J.R., and Callaway, E.M. (2020). Extraction of Distinct Neuronal Cell Types from within a Genetically Continuous Population. Neuron 107, 274–282 e276.

Kole, M.H., Letzkus, J.J., and Stuart, G.J. (2007). Axon initial segment Kv1 channels control axonal action potential waveform and synaptic efficacy. Neuron 55, 633–647.

Korngreen, A., and Sakmann, B. (2000). Voltage-gated K+ channels in layer 5 neocortical pyramidal neurones from young rats: subtypes and gradients. J Physiol 525 Pt 3, 621–639.

Laing, R.J., Turecek, J., Takahata, T., and Olavarria, J.F. (2015). Identification of Eye-Specific Domains and Their Relation to Callosal Connections in Primary Visual Cortex of Long Evans Rats. Cereb Cortex 25, 3314–3329.

Larkum, M.E., Kaiser, K.M., and Sakmann, B. (1999). Calcium electrogenesis in distal apical dendrites of layer 5 pyramidal cells at a critical frequency of back-propagating action potentials. Proc Natl Acad Sci U S A 96, 14600–14604.

Larkum, M.E., Waters, J., Sakmann, B., and Helmchen, F. (2007). Dendritic spikes in apical dendrites of neocortical layer 2/3 pyramidal neurons. J Neurosci 27, 8999–9008.

Lee, K.S., Vandemark, K., Mezey, D., Shultz, N., and Fitzpatrick, D. (2019). Functional Synaptic Architecture of Callosal Inputs in Mouse Primary Visual Cortex. Neuron 101, 421–428 e425.

Lewis, J.W., and Olavarria, J.F. (1995). Two rules for callosal connectivity in striate cortex of the rat. J Comp Neurol 361, 119–137.

Liang, F., Xiong, X.R., Zingg, B., Ji, X.Y., Zhang, L.I., and Tao, H.W. (2015). Sensory Cortical Control of a Visually Induced Arrest Behavior via Corticotectal Projections. Neuron 86, 755–767.

Longordo, F., To, M.S., Ikeda, K., and Stuart, G.J. (2013). Sublinear integration underlies binocular processing in primary visual cortex. Nat Neurosci 16, 714–723.

Lorincz, A., and Nusser, Z. (2008). Cell-type-dependent molecular composition of the axon initial segment. J Neurosci 28, 14329–14340.

Lund, R.D. (1965). Uncrossed Visual Pathways of Hooded and Albino Rats. Science 149, 1506–1507.

Margrie, T.W., Brecht, M., and Sakmann, B. (2002). In vivo, low-resistance, whole-cell recordings from neurons in the anaesthetized and awake mammalian brain. Pflugers Arch 444, 491–498.

Mason, C., and Slavi, N. (2020). Retinal Ganglion Cell Axon Wiring Establishing the Binocular Circuit. Annu Rev Vis Sci 6, 215–236.

Mazurek, M., Kager, M., and Van Hooser, S.D. (2014). Robust quantification of orientation selectivity and direction selectivity. Front Neural Circuits 8, 92.

Metz, A.E., Spruston, N., and Martina, M. (2007). Dendritic D-type potassium currents inhibit the spike afterdepolarization in rat hippocampal CA1 pyramidal neurons. J Physiol 581, 175–187.

Minciacchi, D., and Antonini, A. (1984). Binocularity in the visual cortex of the adult cat does not depend on the integrity of the corpus callosum. Behav Brain Res 13, 183–192.

Mizuno, H., Hirano, T., and Tagawa, Y. (2007). Evidence for activity-dependent cortical wiring: formation of interhemispheric connections in neonatal mouse visual cortex requires projection neuron activity. J Neurosci 27, 6760–6770.

Mountcastle, V.B. (1978). An organizing principle for cerebral function: the unit model and the distributed system. The Mindful Brain, MIT Press, Cambridge, MA.

Olavarria, J., and Van Sluyters, R.C. (1983). Widespread callosal connections in infragranular visual cortex of the rat. Brain Res 279, 233–237.

Olavarria, J.F. (2001). Callosal connections correlate preferentially with ipsilateral cortical domains in cat areas 17 and 18, and with contralateral domains in the 17/18 transition zone. J Comp Neurol 433, 441–457.

Ordemann, G.J., Apgar, C.J., and Brager, D.H. (2019). D-type potassium channels normalize action potential firing between dorsal and ventral CA1 neurons of the mouse hippocampus. J Neurophysiol 121, 983–995.

Pachitariu, M., Stringer, C., Schröder, S., Dipoppa, M., Rossi, L.F., Carandini, M., and Harris, K.D. (2016). Suite2p: beyond 10,000 neurons with standard two-photon microscopy. BioRxiv 061507.

Petreanu, L., Mao, T., Sternson, S.M., and Svoboda, K. (2009). The subcellular organization of neocortical excitatory connections. Nature 457, 1142–1145.

Priebe, N.J., and McGee, A.W. (2014). Mouse vision as a gateway for understanding how experience shapes neural circuits. Front Neural Circuits 8, 123.

Ramachandra, V., Pawlak, V., Wallace, D.J., and Kerr, J.N.D. (2020). Impact of visual callosal pathway is dependent upon ipsilateral thalamus. Nat Commun 11, 1889.

Restani, L., Cerri, C., Pietrasanta, M., Gianfranceschi, L., Maffei, L., and Caleo, M. (2009). Functional masking of deprived eye responses by callosal input during ocular dominance plasticity. Neuron 64, 707–718.

Ringach, D.L., Bredfeldt, C.E., Shapley, R.M., and Hawken, M.J. (2002). Suppression of neural responses to nonoptimal stimuli correlates with tuning selectivity in macaque V1. J Neurophysiol 87, 1018–1027.

Robertson, B., Owen, D., Stow, J., Butler, C., and Newland, C. (1996). Novel effects of dendrotoxin homologues on subtypes of mammalian Kv1 potassium channels expressed in Xenopus oocytes. FEBS Lett 383, 26–30.

Smith, S.L., Smith, I.T., Branco, T., and Hausser, M. (2013). Dendritic spikes enhance stimulus selectivity in cortical neurons in vivo. Nature 503, 115–120.

Stuart, G.J., Dodt, H.U., and Sakmann, B. (1993). Patch-clamp recordings from the soma and dendrites of neurons in brain slices using infrared video microscopy. Pflugers Arch 423, 511–518.

Stuhmer, W., Ruppersberg, J.P., Schroter, K.H., Sakmann, B., Stocker, M., Giese, K.P., Perschke, A., Baumann, A., and Pongs, O. (1989). Molecular basis of functional diversity of voltage-gated potassium channels in mammalian brain. EMBO J 8, 3235–3244.

Tan, A.Y., Brown, B.D., Scholl, B., Mohanty, D., and Priebe, N.J. (2011). Orientation selectivity of synaptic input to neurons in mouse and cat primary visual cortex. J Neurosci 31, 12339–12350.

Tan, L., Tring, E., Ringach, D.L., Zipursky, S.L., and Trachtenberg, J.T. (2020). Vision Changes the Cellular Composition of Binocular Circuitry during the Critical Period. Neuron 108, 735–747 e736.

Tervo, D.G., Hwang, B.Y., Viswanathan, S., Gaj, T., Lavzin, M., Ritola, K.D., Lindo, S., Michael, S., Kuleshova, E., Ojala, D., et al. (2016). A Designer AAV Variant Permits Efficient Retrograde Access to Projection Neurons. Neuron 92, 372–382.

Turrigiano, G.G., and Nelson, S.B. (2004). Homeostatic plasticity in the developing nervous system. Nat Rev Neurosci 5, 97–107.

Weiler, S., Guggiana Nilo, D., Bonhoeffer, T., Hubener, M., Rose, T., and Scheuss, V. (2023). Functional and structural features of L2/3 pyramidal cells continuously covary with pial depth in mouse visual cortex. Cereb Cortex 33, 3715–3733.

Welchman, A.E. (2016). The Human Brain in Depth: How We See in 3D. Annu Rev Vis Sci 2, 345–376.

Yinon, U., Chen, M., and Gelerstein, S. (1992). Binocularity and Excitability Loss in Visual-Cortex Cells of Corpus-Callosum Transected Kittens and Cats. Brain Research Bulletin 29, 541–552.

Zhao, X., Liu, M., and Cang, J. (2013). Sublinear binocular integration preserves orientation selectivity in mouse visual cortex. Nat Commun 4, 2088.

Zingg, B., Chou, X.L., Zhang, Z.G., Mesik, L., Liang, F., Tao, H.W., and Zhang, L.I. (2017). AAV-Mediated Anterograde Transsynaptic Tagging: Mapping Corticocollicular Input-Defined Neural Pathways for Defense Behaviors. Neuron 93, 33–47.

